# Enhanced fluorescence lifetime imaging microscopy denoising via principal component analysis

**DOI:** 10.1101/2025.02.26.640419

**Authors:** Soheil Soltani, Jack G. Paulson, Emma J. Fong, Shannon M. Mumenthaler, Andrea M. Armani

**Affiliations:** Ellison Medical Institute, Los Angeles, California 90064, USA; Mork Family Department of Chemical Engineering and Materials Science, Viterbi School of Engineering, University of Southern California, Los Angeles, California 90089, USA; Keck School of Medicine of USC, University of Southern California, Los Angeles, California 90033, USA; Alfred E. Mann Department of Biomedical Engineering, Viterbi School of Engineering, University of Southern California, Los Angeles, California 90089, USA; Ming Hsieh Department of Electrical and Computer Engineering – Electrophysics, Viterbi School of Engineering, University of Southern California, Los Angeles, California 90089, USA

**Keywords:** Fluorescence Lifetime Microscopy, FLIM, Phasor Analysis, Noise Reduction, Denoising

## Abstract

Fluorescence Lifetime Imaging Microscopy (FLIM) quantifies the autofluorescence lifetime to measure cellular metabolism, therapeutic efficacy, and disease progression. These dynamic processes are intrinsically heterogeneous, increasing the complexity of the signal analysis. Often noise reduction strategies that combine thresholding and non-selective data smoothing filters are applied. These can result in error introduction and data loss. To mitigate these issues, we develop noise-corrected principal component analysis (NC-PCA). This approach isolates the signal of interest by selectively identifying and removing the noise. To validate NC-PCA, a secondary analysis of FLIM images of patient-derived colorectal cancer organoids exposed to a range of therapeutics was performed. First, we demonstrate that NC-PCA decreases the uncertainty up to 4-fold in comparison to conventional analysis with no data loss. Then, using a merged data set, we show that NC-PCA, unlike conventional methods, identifies multiple metabolic states. Thus, NC-PCA provides an enabling tool to advance FLIM analysis across fields.

## 1 Introduction

Fluorescent lifetime imaging microscopy (FLIM) is a versatile imaging method used across cell biology, neuroscience, and cancer research^1–8^. Unlike conventional fluorescent imaging which detects the emission intensity, FLIM analyzes the fluorescent lifetime. In the context of biology, changes in fluorescent lifetimes can be correlated to protein-protein interactions^8–10^, changes in pH or temperature^11–13^, and molecular dynamics^14–18^. Moreover, FLIM enables the detection of the intrinsic autofluorescent lifetime signal that is characteristic of cellular metabolism^19–21^. Given its impact, several approaches for FLIM data analysis have been developed.

Originally, FLIM leveraged exponential fitting methods to extract lifetimes, where a simple linear combination of exponentials was used to fit per-pixel intensity decays^22–25^. The shortcomings and high computational costs associated with fitting have directed the field towards fit-free analysis methods that are easier to interpret, such as phasor analysis. This method translates FLIM data into Fourier space and maps the intensity decay per pixel onto orthogonal vectors (sines and cosines)^26,27^. Any combination of these components corresponds directly to the pixels and represents a unique lifetime combination. This approach can reveal single or multiple dominating lifetimes through clustering in distinct regions on the phasor histogram plot^26,28^. However, transformations into phasor space inherently integrate noise which complicates the identification of distinct clusters and subtle shifts in the phasor distributions^29^.

Common approaches to remedy the sensitivity to noise in phasor analysis include threshold phasor analysis (TPA) and filtered phasor analysis (FPA)^29–31^, as shown in Fig. 1 a-b. While effective for small datasets with low-noise signals, these methods have two main limitations. First, for larger datasets, adaptive thresholding is required to account for varying noise levels. Second, the filters used in FPA are susceptible to introducing smoothing errors^32^. Therefore, frequent filter parameter tuning is required.

**Fig. 1:**
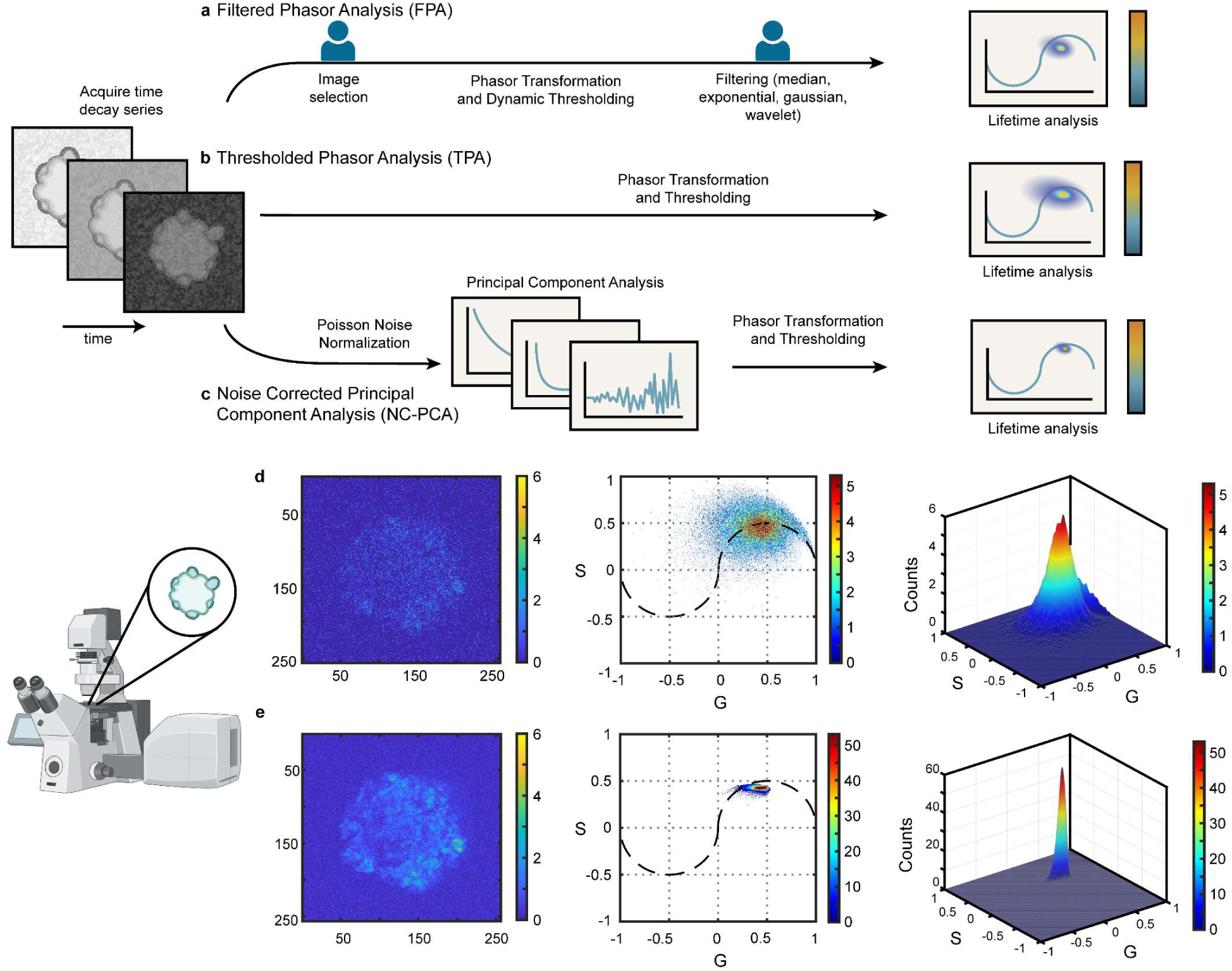
Phasor analysis workflows for FLIM data. After image acquisition on a FLIM system, the data is analyzed using different approaches. (**a**) Filtered phasor analysis method (FPA) includes image selection based on total counts, dynamic thresholding, and application of a selective smoothing filter. (**b**) Thresholded phasor analysis (TPA) only involves the application of an adaptive threshold. (**c**) Noise-corrected Principal Component Analysis (NC-PCA) begins with a Poisson noise normalization, then application of PCA, and subsequent thresholding and phasor transformation. (**d**) Example of results obtained using the TPA analysis method. First time-bin image of a patient-derived colorectal cancer organoid, 2D phasor histogram, and 3D phasor histogram. (**e**) Example of results obtained using the NC-PCA method. Note the reduction in S and G spread in the NC-PCA phasor plot and the 10x difference in counts between parts (**d)** and (**e**) as a result of the improved phasor clustering.

An alternative approach for analyzing large data sets can be found in the statistical analysis method known as Principal Component Analysis (PCA). This technique selectively removes noise while retaining structured data through a transformation into a new orthonormal basis set (Singular Value Decomposition)^33,34^, which is constructed by the eigenvectors and eigenvalues of a mean centered covariance matrix. These vectors, or principal components, are sorted in decreasing variance order. Thus, PCA enables precise identification of important data features with highest variance and facilitates effective removal of noise in complex datasets. Additionally, because it is a data-driven denoising scheme and does not rely on *a priori* knowledge^35^, it removes potential bias. As a result, PCA has been effectively applied in numerous fields, including data compression, omics research, and financial analysis^36–38^.

In the present work, we develop and demonstrate a noise-corrected-PCA (NC-PCA) method for analyzing FLIM data (Fig. 1c). Our NC-PCA analysis workflow denoises time-domain FLIM signals and enhances phasor domain accuracy. The shot noise is reduced, and the correlated linearity of corresponding pixels is preserved^36,39^. To validate the method, we created and analyzed synthetic FLIM data as well as a standard fluorescent dye. Next, we used the NC-PCA method to analyze patient-derived colorectal cancer organoids and confirmed the improvement in data retention and signal integrity as compared to the TPA approach. Finally, we analyzed the metabolic activity of patient-derived colorectal cancer organoids in response to standard therapeutics using all three approaches outlined in Fig. 1. In comparison to TPA and FPA, NC-PCA demonstrated a reduction in error as high as ∼58% and ∼45%, respectively. Finally, we demonstrated that NC-PCA can reveal multiple lifetimes present in systems with untreated and treated organoids superimposed on the same phasor histogram, setting the stage for the discovery of weak biological responses^5,40^.

## 2 Results

### 2.1 Validation of NC-PCA using synthetic data

To demonstrate the effectiveness of NC-PCA for noise reduction in FLIM data, we created a synthetic time-series dataset based on a cell image with geometrically distinct features representing cellular structures (Fig. 2a). The total counts were varied from 200 to 1000 photons to reflect experimental conditions.

**Fig. 2:**
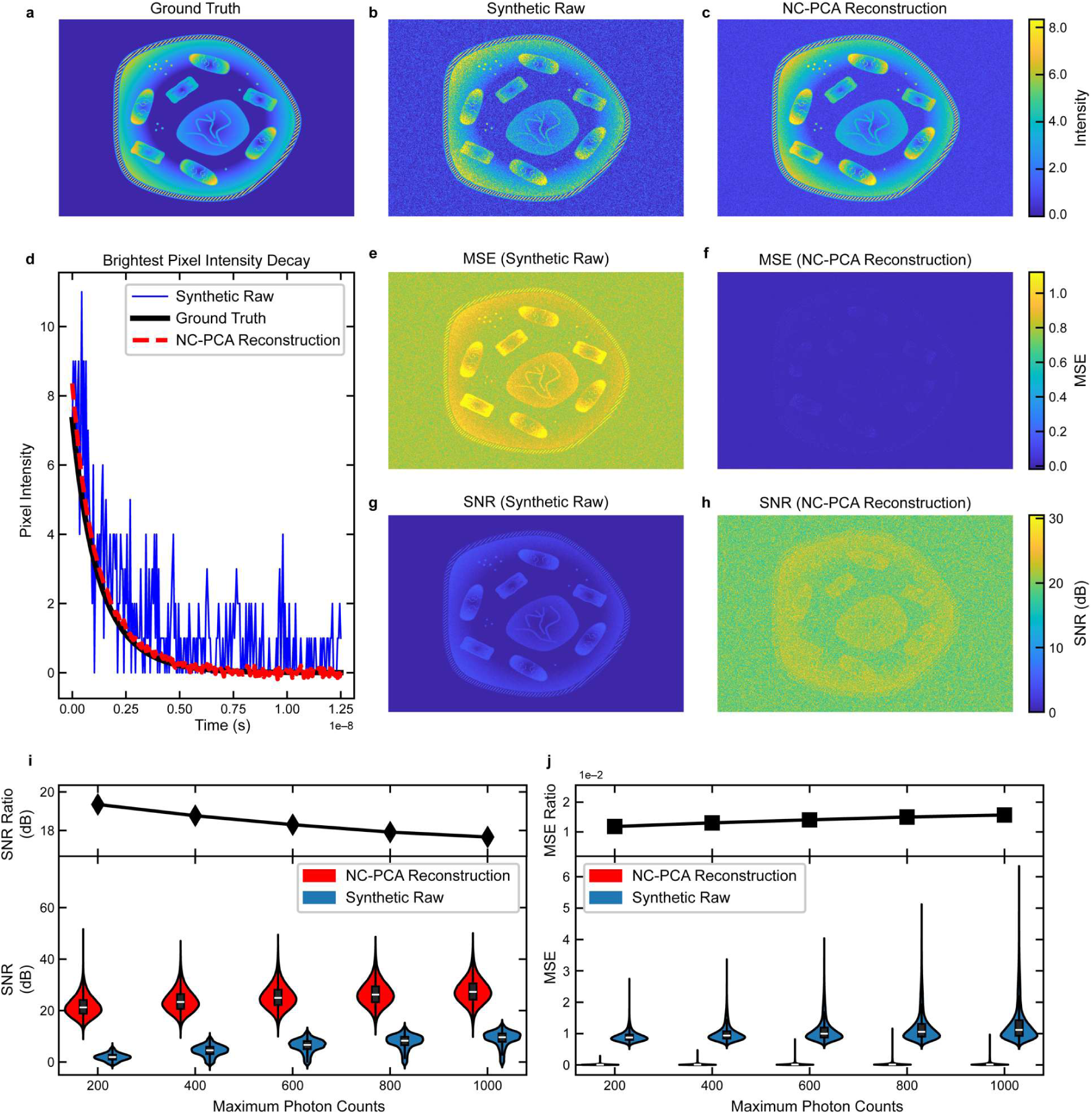
Ground truth comparison of NC-PCA on synthetic FLIM data. The first time bin of the synthetically created data that serves as the (**a**) ground truth and(**b**) that is modified with Poisson noise to serve as the synthetic raw data. A range of photon count levels were investigated, and this data is limited to 200 photon counts, representing extremely noisy data. (**c**) The image shown in part **(b)** is reconstructed using NC-PCA. (**d)** The intensity of the brightest pixel in the ground truth image (solid black), synthetic raw data (solid blue), the NC-PCA (red dashed). (**e, f**) MSE and (**g, h**) SNR are calculated for each pixel across the entire synthetic raw image or NC-PCA reconstructed image. (**i, j**) To explore the role of noise in reconstruction ability, the photon counts were varied from 80 photon counts (low SNR) to 1800 photon counts (High SNR). All results are included in the SI, and a subset of the results are presented. The violin plots for (**i)** SNR and **(j**) MSE images from synthetic raw and NC-PCA reconstruction images highlighting the improvement offered by NC-PCA, in terms of data spread and relative SNR and MSE values. NC-PCA improvement was quantified via SNR and MSE Ratios between synthetic raw and NC-PCA reconstruction.

Fig. 2b-c are the first time bins of the synthetic FLIM dataset and NC-PCA reconstruction with a total photon count of 200. The time dependence of the highest intensity pixel is plotted in Fig. 2 d, and the signal to noise ratio (SNR) and mean square error (MSE) relative to the ground truth were calculated to analyze the quality of the reconstruction. The highest intensity pixel’s SNR increased nearly 3x, from 5.99 dB to 17.5dB, while the MSE decreased from 1.34 to 0.05, highlighting NC-PCA’s ability to enhance signal fidelity.

The SNR and MSE was analyzed for the entire data set of 388,800 pixels over 256 time bins, and Fig. 2e-h present spatial distribution. Notably, the scale bars are unified in these figures to allow direct comparison. The median SNR improves from 1.87 dB to 21.2 dB and the MSE reduces from 0.875 to 0.0101. Overall, NC-PCA yields a ∼20 dB improvement in SNR and ∼90x reduction in MSE.

To further explore the potential of NC-PCA in FLIM analysis, we evaluated the improvement in SNR and MSE as photon counts are increased from 200 to 1000. The distributions of SNR and MSE at each photon count level are shown in the violin plots in Fig. 2i-j. As shown in Fig. 2i-j, NC-PCA improved both SNR and MSE across the entire range studied. These findings underscore the ability of NC-PCA to enhance signal fidelity specially in low photon count pixels. Additional analysis over an extended photon count range and the corresponding images are in the SI.

In addition, we define two metrics to quantify the level of improvement NC-PCA achieves: the SNR Ratio, which compares the difference between SNR medians in log space per-distribution, and MSE Ratio, which is the ratio between MSE medians per-distribution. These two metrics further quantify NC-PCA’s effectiveness. As shown in Fig. 2i-j, both ratios favored the NC-PCA method. As photon counts increase, the SNR of the synthetic image increases, allowing the decay signal per pixel to approach the ground truth signal. Consequently, while an overall improvement in SNR and MSE persists, the magnitude of this improvement diminishes at higher photon counts.

To validate the NC-PCA method experimentally, Coumarin-6, a standard fluorophore with a known lifetime of 2.43-2.60^41–43^, was imaged using 30- and 100-frame acquisitions. The 100-frame measurement was used as ground truth and 30-frame as noisy data. Next, we applied a single component exponential fit to every pixel in each image to calculate lifetime values. In addition, NC-PCA was performed on the 30-frame acquisition data, and lifetime values were obtained. However, due to high noise, we were unable to obtain a lifetime value from the 30-frame results without application of NC-PCA. All data is summarized in the SI (Fig. S9). Based on these results, the ground truth histogram has a mean of 2.58 ns and a Full Width Half Maximum (FWHM) of 0.2 ns, in agreement with prior results^41–43^, while the 30-frame NC-PCA data shows a mean of 2.56 ns and a FWHM of 0.23 ns. This demonstrates that NC-PCA enhances low-frame data to match high-frame ground truth, enabling retrieval of lifetime information reliably.

### 2.2 Validation using patient-derived colorectal cancer organoids

FLIM imaging is routinely used to evaluate a biological system’s response to a perturbation by performing a phasor transformation of FLIM data and analyzing two variables, G and S. Together these parameters represent the position and direction of shifts on the phasor histogram. The shifts in position and direction reveal information regarding the fluorescent lifetime of different components in a sample, which can be correlated to changes in metabolic activity^19,20,40^.

When a single lifetime is present, G and S fall on the universal unit semicircle. Shorter lifetimes are located closer to the edge of the semicircle, and longer lifetimes are closer to the origin. For any other points inside the semicircle, lifetime is defined as the weighted sum of the lifetimes on the semicircle^27^. Therefore, the position of the phasor histogram determines the exact composition of lifetimes. However, the noise content of the decay signal in every pixel broadens the phasor histogram and increases measurement error^25,44^.

The ability of NC-PCA to overcome this limitation is demonstrated using FLIM images of patient-derived colorectal cancer organoids. Similar to the previous measurement, we used 100-frame data as the reference data and 10-frame imaging with and without NC-PCA for the experimental measurement. The first time-bin images for all three measurements are shown in Fig. 3a-c, and the associated phasor plots are in Fig. 3d-f.

**Fig. 3:**
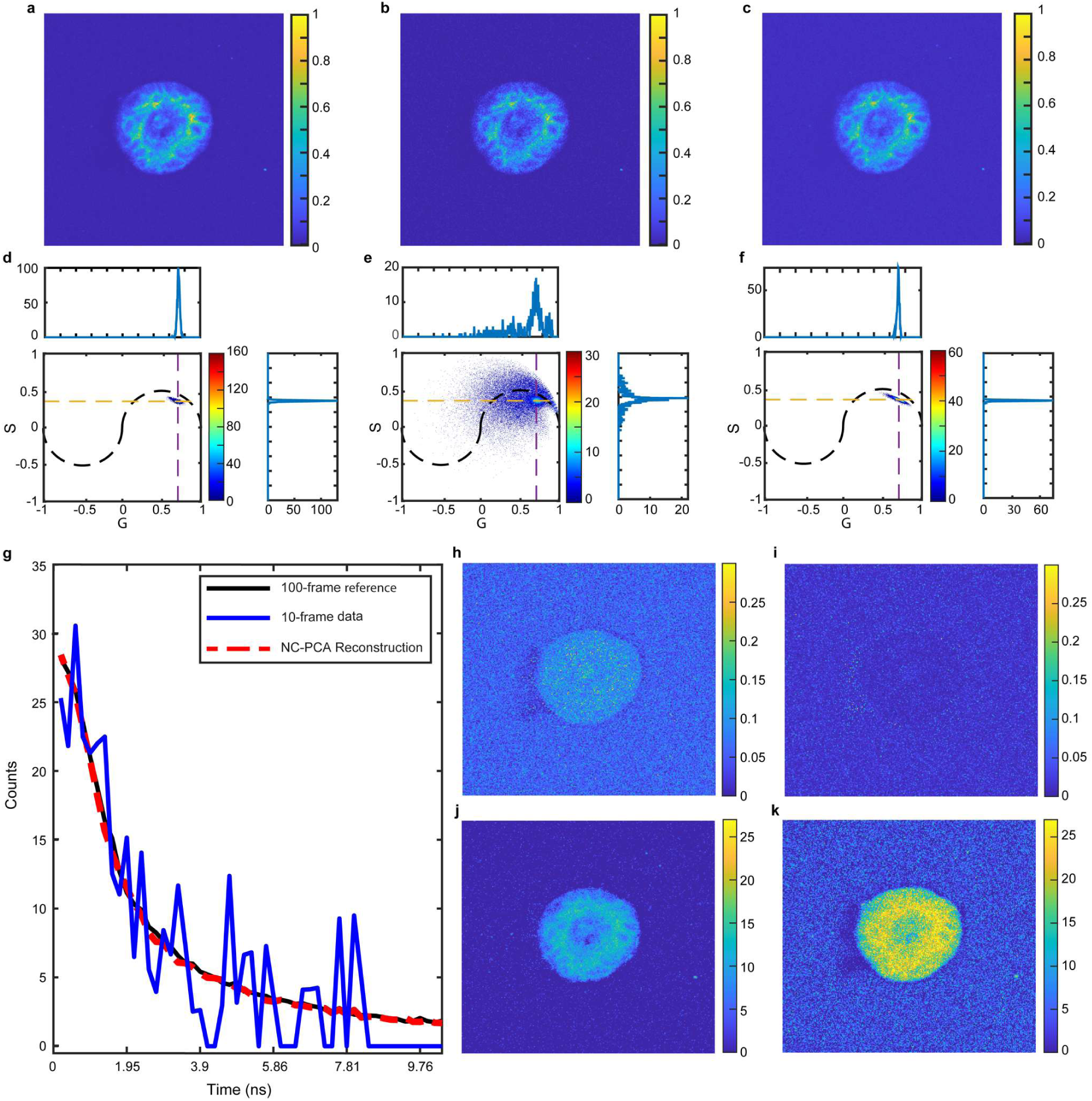
Application of NC-PCA on a patient-derived colorectal cancer organoid. FLIM images of (**a**) 100-frame FLIM data used as low-noise reference data, (**b**) 10-frame FLIM data, and (**c**) NC-PCA denoised 10-frame FLIM data. (**d**-**f**) Phasor plot of (**d**) reference data, (**e**) 10-frame FLIM data, and (**f**) NC-PCA denoised 10-frame FLIM data. The median count coordinates and the FWHM along the G and S axes are calculated along the dashed lines (**g**) The intensity of the brightest pixel in the reference image (solid black), 10-frame image (solid blue), and NC-reconstruction (red dashed) tracked over time. (**h, i**) MSE and (**j, k**) SNR are calculated for each pixel across the entire (**h, j**) 10-frame image and (**i, k**) NC-PCA reconstructed image using the 100-frame data set as the reference.

Qualitatively, the G and S median values appear similar for all three plots; however, the distributions vary greatly. To quantify the difference, the median count coordinates and the FWHM along the G and S axes are calculated along the dashed lines in Fig. 3d-f. The calculated values for G and S median coordinates for the NC-PCA denoised 10-frame data and the 100-frame reference data differ by less than 0.3% and 0.25%, respectively. This value is within the error of the measurement and demonstrates that NC-PCA does not alter the absolute value of the result. In contrast, the FWHM of the NC-PCA reconstruction along either the G or S axis is nearly 5x narrower than the initial 10-frame data set, confirming the performance improvement.

The reconstruction quality was further quantified by calculating the SNR and MSE using the 100-frame data set as the reference data. For the given pixel, the SNR increased by 9.7dB, and the MSE decreases by ∼9.4 when the NC-PCA method is applied (Fig. 3g). When the entire data set is evaluated, similar trends in SNR (Fig. 3h-i) and MSE (Fig. 3j-k) are observed. We find an average improvement in the SNR by ∼10 dB and a decrease in MSE by ∼10 fold.

Additionally, NC-PCA reconstruction adaptively denoises the low-frame FLIM signal, resulting in an improvement in the time domain data and consequently, in the fit-free phasor domains. This improvement significantly enhances the phasor domain detection accuracy with no information lost during the process. The phasor domain detection resolution improved from 0.276 to 0.052 as the frame number decreased from 100-to 10-frames.

### 2.3 Assessment of Data Loss

FLIM can detect metabolic changes in biological systems by monitoring the change in auto-fluorescent behavior of NADH as it shifts between free and protein-bound states^21,40,45,46^. Because these two states have distinctly different lifetimes, population changes are evident on the phasor histogram. Typically, the results are reported in terms of the fraction bound (fB) NADH, which is directly proportional to G and S (see SI).

Like many biological imaging measurements, FLIM data can suffer from significant noise that is integrated directly into G and S. To simplify the data interpretation and increase statistical confidence, the phasor distribution is often collapsed into a single pair of S and G values post-filtering. This technique requires that the entire phasor distribution of each image is averaged, and the histogram is then approximated by a single pair of values. This simplification may lead to information loss. By reducing signal noise, while preserving lifetime dynamics, NC-PCA offers a potential solution to mitigate this trade-off.

To demonstrate the capability of NC-PCA to obviate the need for single-point approximation, we calculated the phasor histograms of a representative image and the associated quantitative information loss metrics. Fig. 4a-c presents the phasor histogram, G, and fB images, respectively, of the FLIM data processed using the FPA process. Fig. 4d-f shows the same image processed using the NC-PCA method.

**Fig. 4:**
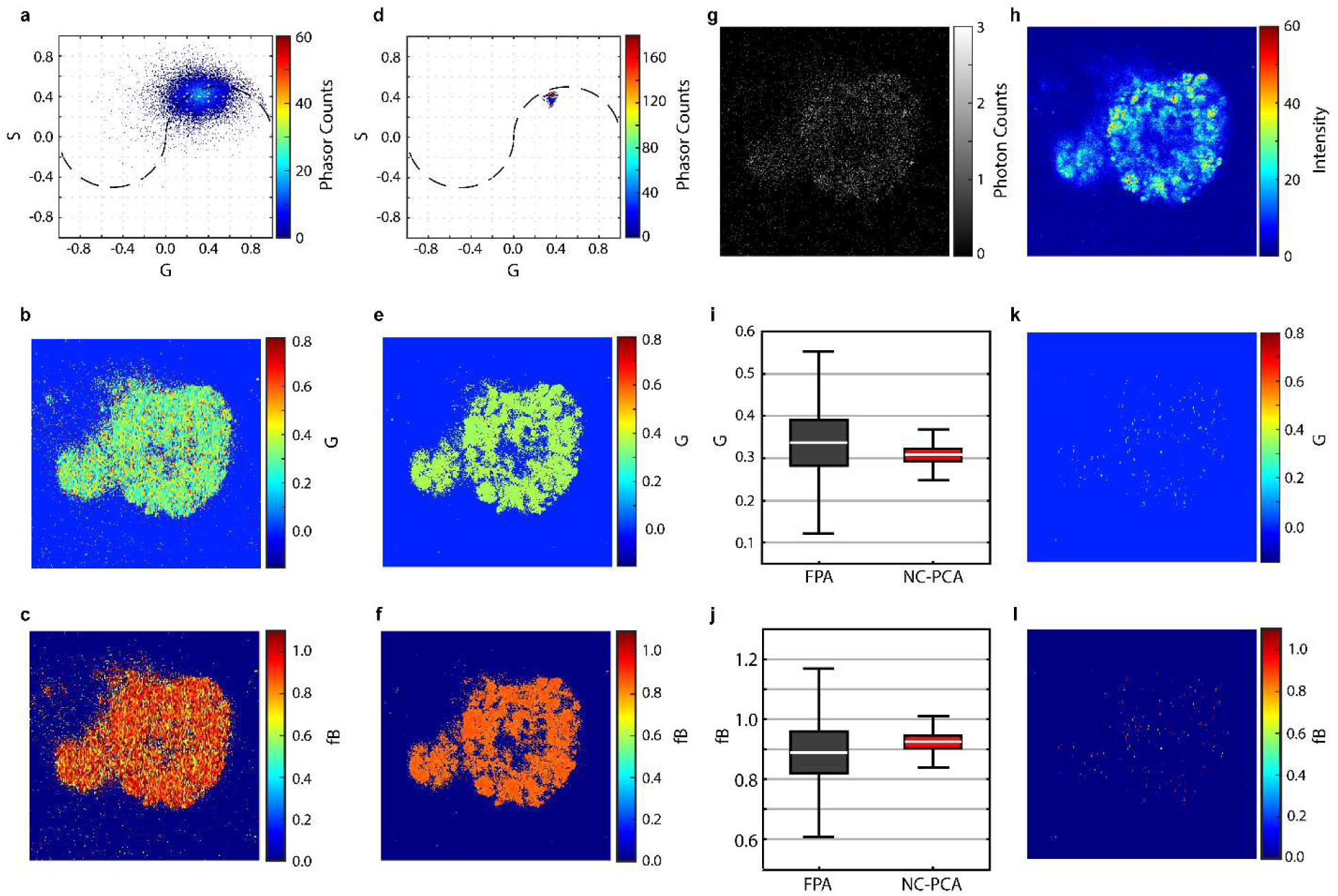
Comparison of phasor distributions of FLIM images of a patient-derived colorectal cancer organoid using FPA and NC-PCA analysis. Phasor histogram, G image, and fraction bound (fB) image of a organoid created using (**a-c**) FPA method or (**d-f**) NC-PCA method. Data was taken using 10-frame acquisition. (**g**) Greyscale image of the first time bin. (**h**) Total photon count per pixel of the same FLIM image used for the phasor histogram in (**a**). (**i, j**) Comparison of boxplots for G and fB distributions when the data is analyzed using either FPA or NC-PCA method. (**k, l**) The pixel maps of G and fB images, highlighting the degree of data loss that occurs during the FPA process if averaging is applied. The pixels that preserve their original values if the entire phasor histogram is approximated by average G and S values (data preservation map).

While applying a median filter helps localize the phasor histogram and smooths the G and fB images, the resulting distributions still display significant spread (Fig. 4a-c). This ambiguity necessitates the use of aggressive thresholding, filtering, and subsequent phasor averaging. In contrast, the NC-PCA method effectively denoises the phasor histogram, as well as the G and fB images, reducing the uncertainty and spread of phasor distribution (Fig. 4d-f). Therefore NC-PCA precludes the need for post-filter averaging.

The effective denoising capabilities of NC-PCA is further illustrated when the images obtained using both methods are compared. The images from filtered phasor analysis (Fig. 4b-c) closely resemble the first time-bin image which is a noisy representative of the FLIM image (Fig. 4g). In contrast, the G and fB images derived from NC-PCA (Fig. 4e-f) more closely correspond to the total count map (Fig. 4h), an aggressively filtered representation of the FLIM data. The reduction in the phasor distribution using NC-PCA is quantitatively illustrated in Fig. 4i-j. There is a ∼4-fold improvement between the FWHM of the phasor and fB distributions using NC-PCA compared to FPA method.

To show the degree of information loss incurred through the FPA process, we calculated the map of the G and fB values that maintain their original values when the phasor histogram is averaged and approximated by a single pair of G and S. The results are shown in Fig. 4k-l. Less than 1% of data points in G and fB maps are preserved during the FPA process, emphasizing that averaging the FLIM histogram can result in significant information loss. In contrast, NC-PCA denoises the FLIM data obviating the need for an averaging step and mitigating this information loss.

### 2.4 Revealing Metabolic Dynamics

In the context of FLIM analysis, the minimum resolvable signal change on the phasor histogram is the FWHM or signal linewidth of the phasor accumulation point. Broad phasor distributions and associated histogram linewidths limit the ability to detect subtle metabolic responses. To demonstrate the impact of NC-PCA in overcoming this challenge, we conducted a secondary data analysis on an existing dataset of patient-derived colorectal cancer organoids treated with a range of therapeutics (Staurosporine (ST), 5-flourouracil, 7-ethyl-10-hydroxycamtothecin, Cetuximab, and 3-bromopyruvate) and range of doses (0.01 to 100 μM for 72 hours)^40^.

In the first study, the phasor distributions from 10 data sets of an untreated and 72-hr ST-treated organoids are calculated and then merged, creating an overlayed data set that contains two known metabolic states. Fig. 5a-d shows representative first time-bin images and phasor histograms for an untreated organoid processed using TPA and NC-PCA methods. Due to the irreversibility in the transformation into the phasor domain, FPA is unable to reconstruct the image and therefore is not shown. Phasor plots of the whole dataset were analyzed (Fig. 5e-f, h-i) and overlaid (Fig. 5g, j). In parallel, the histogram of the data along the G axis was calculated and is plotted as part of Fig. 5e-j.

**Fig. 5:**
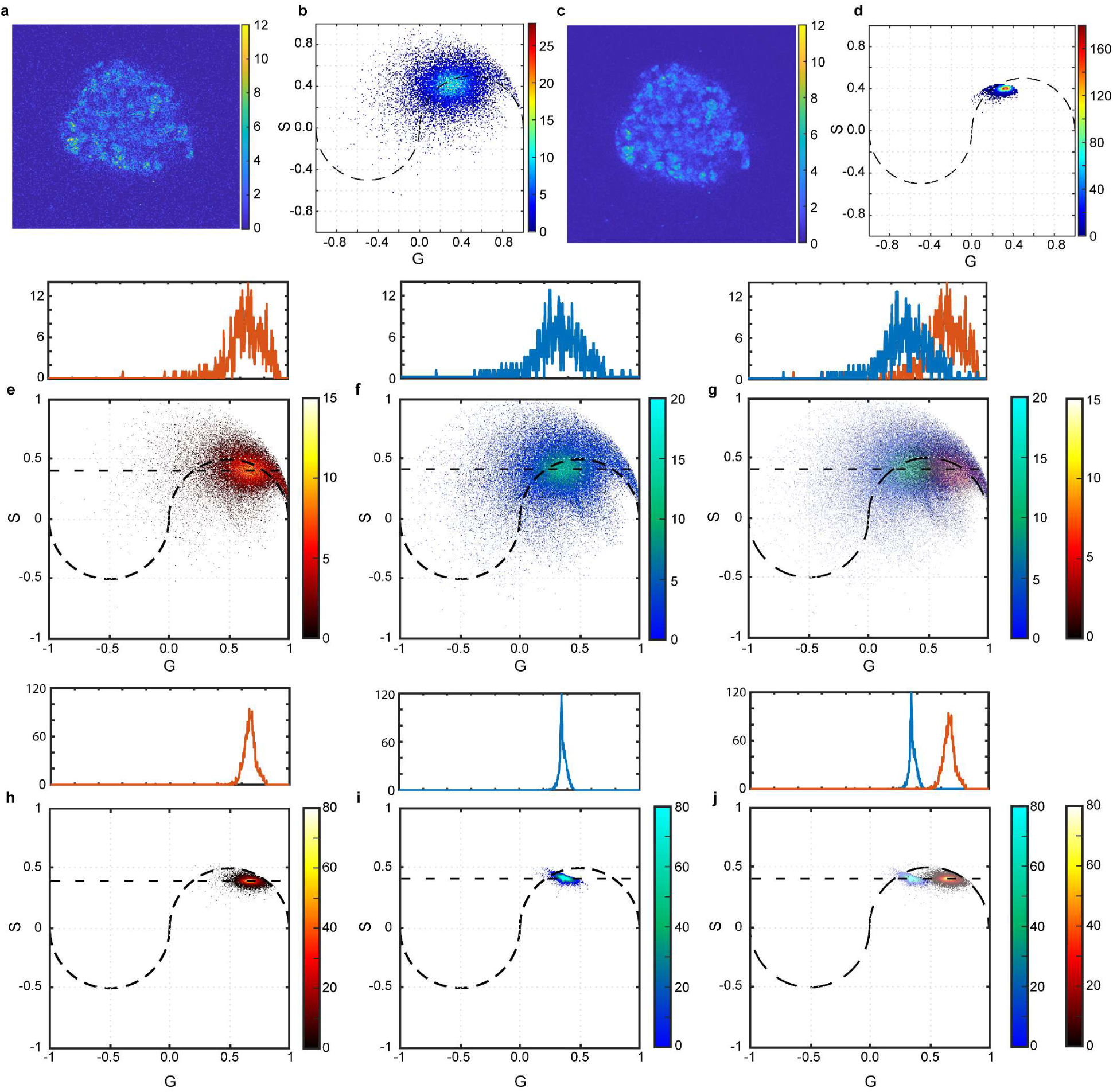
Detecting time dependent effects of ST on patient-derived colorectal cancer organoid metabolism. First time bin and corresponding phasor histogram of an untreated patient-derived colorectal cancer organoid data analyzed using (**a, b**) TPA approach or (**c, d**) NC-PCA approach. Data was taken using 10-frame acquisition. (**e**) Phasor histograms of untreated and (**f**) 72 hour ST-treated organoids analyzed using the FPA approach. (**g**) The phasor histograms in parts (**e**) and (**f**) are overlaid. The count distributions along the maximum count lines are fit to Gaussian distributions. (**h, i**) The same data set is analyzed using the NC-PCA approach, and (**j**) the pair of phasor histograms are overlaid. The count distributions along the maximum count lines are fit to Gaussian distributions.

As evident in Fig. 5e-f, the two phasor distributions generated using FPA have significant overlap. Based on the FWHM of the associated histograms, the minimum detectable signal change on the histogram is limited to 0.5446. In Fig. 5g, the separation distance between these histograms, defined as the difference in G-coordinates of their maximum counts, is ∼0.31. This separation is nearly half the FWHM value. This analysis highlights why multiple metabolic states or metabolic heterogeneity within a single frame are not distinguishable when FPA is used. This limitation can be directly attributed to the denoising approach used in FPA.

In contrast, NC-PCA effectively denoises the FLIM data and reduces the phasor histogram FWHM, resulting in clear identification of the two metabolic states (Figure 5h-j). Specifically, the signal linewidth is 0.105, which is improved by ∼5.5 times in comparison to the FPA method. The separation between the two states on the phasor histogram is ∼0.31, which is ∼3 times larger than the phasor histogram linewidth, making identification of sample heterogeneity straightforward. Importantly, the differences in the mean values of G as determined by both methods are not statistically significant, indicating that the NC-PCA method is not altering the data. Thus, NC-PCA can reveal multiple metabolic processes happening within a biological system that are not clearly distinguishable by conventional methods. Furthermore, NC-PCA can identify these distinct characteristics without the express need for extra smoothing or complex wavelet filtering.

In the second study, the boxplots of G and fB values were calculated for the entire dataset across ST concentrations and control samples using TPA, FPA, and NC-PCA. Additional analysis for varying concentrations and all alternative drug treatments are in the SI. As seen in Fig. 6a-b, NC-PCA can correct skewed or spread phasors outside physically relevant states, which occur with TPA and FPA due to noise. Values outside the universal semicircle or fraction bound values above 1 or below 0 are physically impossible. NC-PCA corrects these tendencies, restoring biologically relevant values.

**Fig. 6:**
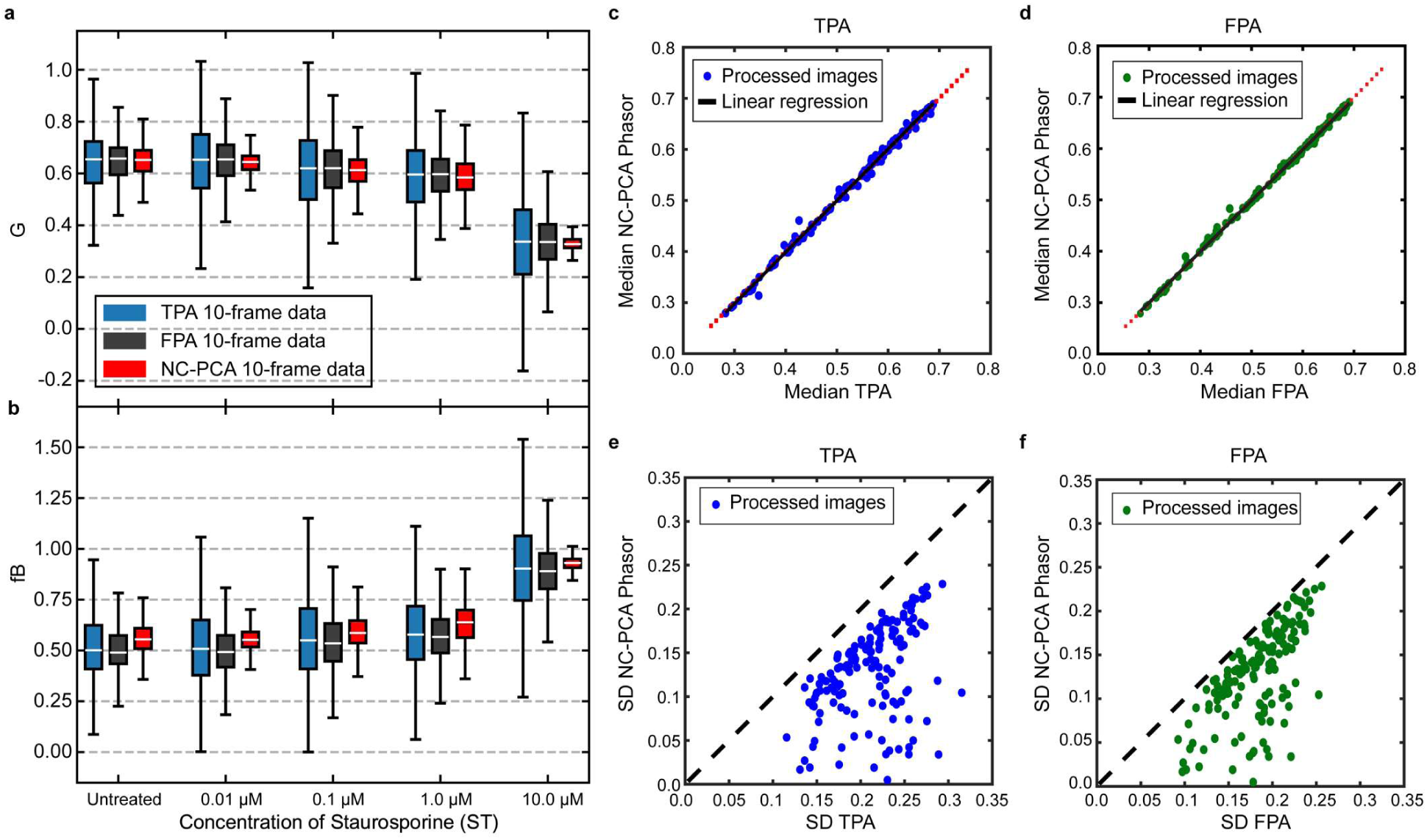
Ability of NC-PCA to improve accuracy of ST dose-response on phasor histogram. (**a**) Boxplot of G component of phasors and (**b**) fB values analyzed using TPA (blue), FPA (grey), and NC-PCA (red) for untreated, 0.01, 0.1, 1, and 10.0 μM of ST. (**c, d**) Comparison of median G values for NC-PCA vs FPA (blue circles) and NCA-PCA vs TPA (green circles). As a reference, the line y=x is shown (red dash). For both data sets, the majority of the data lies on the y=x line. The calculated slope is 1.0097 and 0.998 for NC-PCA relative to TPA and to FPA (black line). (**e, f**) Comparison of standard deviation for G values between NC-PCA vs FPA (blue circles) and NCA-PCA vs TPA (green circles). As a reference, the line y=x is shown (light gray dash). Across all data sets, the standard deviation decreases when NC-PCA is used in place of either TPA or FPA.

A key aspect of any denoising or filtering method is that it should preserve the information and the structure of the data. Based on the prior measurements, we expect that NC-PCA should have a negligible effect on the median values for single cluster phasor distributions and simultaneously reduce the noise-associated spread. In Fig. 6a, it appears that NC-PCA does not significantly change the median values in G. To quantitatively determine if the data median is preserved, we created a single pooled data set (size n=150 images), analyze it using TPA, FPA and NC-PCA phasor analysis methods, and evaluate the median values of G (Fig. 6c-d). As can be observed in Fig. 6c-d, the median value across all experimental conditions is independent of data analysis method, confirming that the data is preserved.

The primary objective of NC-PCA is to reduce error. Fig. 6a-b shows a reduction in the upper and lower quartiles and whisker lengths across all ST concentrations with NC-PCA. To quantify this, the standard deviation (SD) of G for the pooled dataset was calculated. Fig. 6e-f demonstrates a notable SD reduction with NC-PCA, indicating consistent phasor distribution dynamics while effectively removing noise, and preserving key information.

Comparing SD values in Fig. 6e-f highlights an interesting feature of NC-PCA. FPA results in smaller SD values than TPA, yet NC-PCA achieves the lowest SD among methods. This demonstrates that while traditional filters remove noise to some extent, their effectiveness is limited compared to NC-PCA. Unlike conventional methods, NC-PCA efficiently denoises data without the extreme parameter tuning necessary in the TPA and FPA methods. A more comprehensive analysis across different FLIM systems, additional therapeutics, and doses are presented in the SI.

## 3 Discussion

A crucial aspect of measuring the metabolic activity of any biological system using FLIM is reducing intrinsic noise in the data while preserving signal integrity. The rapid data acquisition required for metabolic studies results in inherently low photon counts and low SNR, making this goal particularly challenging. The NC-PCA-based FLIM image analysis method addresses this challenge by preserving signal intensity and decay structure while selectively isolating noise, thereby increasing SNR. As a result, NC-PCA enables the effective application of advanced techniques such as reference-free calibration, multi-exponential lifetime fitting, fluorescence lifetime-resolved anisotropy, and machine learning-based FLIM analysis^10,17,20,29,41^.

In addition to improving data analysis accuracy, NC-PCA efficiently handles large FLIM datasets with minimal manual intervention. This reduces the need for extensive parameter tuning, streamlining analysis for high-throughput studies while enhancing reproducibility and reliability in large-scale research. Taken together, these advantages position NC-PCA as a powerful and enabling tool for both fundamental biological discovery and translational research. For example, its higher precision can reveal multiple metabolic states within the same sample, providing deeper insights into biological heterogeneity. Additionally, it improves our ability to decipher the response dynamics and efficacy of therapeutics, ultimately advancing biomedical research^17,20,31,40,47,48^.

## 4 Online Methods

### 4.1 Data Analysis Methods

The FLIM data was analyzed using TPA, FPA, and NC-PCA through custom Python and MATLAB scripts. Notably, both scripts operate independently and produce identical results. Both the Python and MATLAB scripts are freely available on Zenodo (https://doi.org/10.5281/zenodo.14895223). While both scripts primarily process raw time-series TIFF files, they are designed for customization and adaptability. The algorithm implementation in both Python and MATLAB consists of three main components: a data preprocessing script, an NC-PCA analysis script, and a phasor analysis script. Adapting non-TIFF file formats for compatibility with the NC-PCA and phasor analysis scripts simply requires converting the imported data into arrays.

The TPA workflow is the simplest of the three. This method takes time-series FLIM data and applies an intensity (integrated counts) threshold and processes the data through phasor analysis. Intensity thresholding is performed by summing the total counts per pixel over time, applying a minimum count threshold, and setting all pixel values at or below that threshold to zero across each time bin.

The FPA workflow incorporates both intensity thresholding (as defined above) and median filtering. The median filter is applied to the phasor histogram, a common practice in similar applications. A kernel (or window size) is specified, determining the number of local data points used for filtering. While intensity thresholding is performed before any phasor transformation, the median filter is applied in phasor space and is included as a functionality in both the Python and MATLAB phasor analysis scripts.

The NC-PCA workflow begins by defining the number of principal components for projecting the raw data. If this number is unknown, the Python script includes a selection process using a scree plot, which visualizes the total variance captured by each principal component. For this study, we found that three principal components optimally preserved signal while removing noise. The script then applies the noise-correction (NC) scheme and stores the NC factors for later use. The Python script also allows tunability in the PCA method selection, depending on dataset size and computational efficiency. For this research, we used Singular Value Decomposition (SVD) to extract the principal components. Once the dataset was fitted and the principal components were extracted, the original data was projected onto the new basis. The NC factors are then factored into the transformed dataset to properly scale the data back into its original state. Finally, the same intensity thresholding used in TPA and FPA was applied to ensure consistency across all methods. More information on the data analysis methods can be found in the SI.

### 4.2 Creation of Synthetic FLIM data

A series of synthetic images were first created in PowerPoint. Three distinct images were designed to contain: (1) mitochondria and ribosomes, (2) the nucleus and vacuoles, and (3) the cell wall. When combined, these images mimicked a basic cell structure with distinct geometric features. By initially keeping them separate, we could independently tune the lifetime characteristics associated with each component set. Each image was assigned a lifetime and projected into 256 time bins, with signals decaying exponentially according to their specified lifetimes. Once the temporal behavior was modeled, the three time-series datasets were merged into a single dataset.

To ensure that noise in the synthetic data accurately matched experimental data, we applied a Poisson noise distribution. Commonly known as Shot noise, the dominant noise source in FLIM data arises from photon counting errors, with a standard deviation proportional to the square root of photon counts. Importantly, time-correlated single-photon detectors count photons independently of time bins. As a result, shot noise is temporally uncorrelated, with dependencies only arising from the signal’s exponential decay. To simulate this noise profile, we applied a randomized Poisson distribution to each pixel^39^, setting the mean value to match the corresponding time-bin photon count. Additionally, to prevent zero standard deviation and ensure nonzero noise levels, a mean value of 0.8 (*λ*=0.8 for Poisson distribution) was assigned to background pixels with no fluorescent lifetime. This value was selected based on experimental results (see SI).

Since the signal-to-noise ratio (SNR) of shot noise scales with the square root of photon counts, we varied noise characteristics by adjusting the total photon count per pixel. The total counts per image were chosen to reflect experimental datasets, ranging from 80 to 1,800 photons. This approach ensured that our calculated noise accurately modeled variations encountered in experimental FLIM data while also testing the upper and lower limits of NC-PCA. Additional details on synthetic FLIM dataset construction are provided in the SI.

### 4.3 Sample preparation protocols

#### Coumarin-6 dye

Coumarin-6 (Sigma Aldrich #442631) reconstituted in 200 proof ethanol was used as the calibration standard before each imaging session.

#### Organoid culture

Patient- derived tumor organoids (PDTOs) were generated from colorectal cancer (CRC) tumor resections received from the USC Norris Comprehensive Cancer Center Translational Pathology Core according to Institutional Review Board (Protocol HS-06-00678) approval. Tissues were processed as outlined previously^49,50^. PDTOs were grown at 37^0^C and 5% CO_2_ in CRC media which contained supplements as outlined in a previous publication^40^ (Table 1, CRC media).

Sample preparation for FLIM imaging was also done according to previous publications. Briefly, PDTOs were resuspended in Reduced Growth Factor Basement Membrane Extract (BME; Trevigen, Gaithersburg, MD; 3533-001-02) at 100,000 cells/mL and 10uL of this suspension was seeded in a 96-well glass-bottom plate (Mattek, Ashland, MA; P96G-1.5-5-F). Plates were allowed to slightly gel at room temperature for 2 minutes and then inverted and incubated at 37^0^C for an additional 10 minutes to allow full gelling of the BME.

PDTOs were allowed to grow for 7 days in 200 μL CRC media before switched out with media spiked with drugs. Cells were dosed with various concentrations of staurosporine (ST Sigma-Aldrich, St. Louis, MO; 569396), 5-fluorouracil (5-FU; Selleck Chemicals, S1209), 7-ethyl-10-hydroxycamtothecin (SN-38; Sigma-Aldrich, St. Louis, MO; H0165), 3-bromopyruvate (3-BP; Sigma-Aldrich, St. Louis, MO; 16490), or cetuximab (CTX, MedChemExpress, Monmouth Junction, NJ; HY-P9905). Untreated controls for STA, SN-38 and 5-FU were wells of 0.01% DMSO in CRC media.

### 4.4 FLIM imaging protocols

For all the data presented in the main text, an Olympus FV3000 system equipped with an A320 FastFLIM FLIMbox (ISS Inc., Champaign, IL), hybrid PMT external detectors (Hamamatsu, Shizuoka, Japan), and a Ti:Sapphire 2-photon laser (Spectra-Physics, Mountain View, CA) was used. To confirm the analysis approach, additional data using different imaging systems were also used. Table S1 lists all systems and settings and the corresponding data sets. Additional experimental details for the imaging results are located in the SI.

For Coumarin-6 measurements, we used a 100-frame FLIM image taken as high signal to noise ratio reference data acting as the ground truth. To acquire the low signal to noise ratio Coumarin-6 image, we used 30-frame averaging. Next for both the reference data and raw data we performed a thresholding step to remove pixels with low counts which was followed by a per pixel single exponential fitting. Next the low-frame data (30-frame) was processed using NC-PCA method and a per pixel exponential fitting was performed on the denoised pixels.

For every patient derived organoid presented in the main text, three z-slices with a step-size of 10 μm from the center (20 μm total imaged z-area) were imaged with a UPLXAPO 20X/0.8NA Olympus objective and excited at 740 nm at 15-20 mW. Emission signal was passed through a 458/64 dichroic for NAD(P)H collection and data collection was done with VistaVision software (ISS Inc., Champaign, IL). FLIM imaging of the organoids was performed after 6 and 72 hours respectively. The images used as reference data were taken at 100-frame averaging and the raw images used for denoising were taken at 10-frame averaging.

The PDTO FLIM dataset in Fig.5 and Fig.6 were originally published in our previous work and is used here for secondary analysis^40^.

## Acknowledgments

We would like to thank the members of the Ellison Medical Institute for scientific discussions, and a special thanks to the Cell Line Team (Michael Doche, Roy Lau, Scott Valena and Pratiksha Kshetri) for cell culture assistance. We would like to thank Dr. Heinz-Josef Lenz and his lab members at the USC Keck School of Medicine for providing access to colorectal cancer patient tissues. We would also like to acknowledge the USC Norris Comprehensive Cancer Center Translational Pathology Core and the Molecular Genomics Core for the collection of patient tissue samples and targeted sequencing of our patient organoid lines (Norris Comprehensive Cancer Center CCSG grant, P30CA014089). We acknowledge financial support from the Office of Naval Research (N00014-24-1-2296) and the National Science Foundation (DBI-2414158) awarded to AMA and from the National Cancer Institute Cancer Systems Biology Consortium (U01CA232137) awarded to SMM.

