## Supplementary info for "Enhanced fluorescence lifetime imaging microscopy denoising via principal component analysis"

### 1 Principal Component Analysis (PCA)

#### 1.1 Overview

Principal Component Analysis (PCA) is an adaptation of Singular Value Decomposition that factorizes a data matrix into three primary vectors and ranks these vectors based on their contributions to the variance of the system. In essence, PCA creates a hierarchical coordinate system based directly on the data that is being decomposed. This allows for a reduction of dimensionality while maintaining the primary characteristics of the original data set. The factorization of matrix, X, is conventionally written as:

$$X = U\Sigma V^T \tag{1}$$

where U is the left singular matrix containing the eigenvectors of $XX^T$, $\Sigma$ is a diagonal matrix of singular values of $XX^T$, and $V^T$ is a matrix comprised of right singular vectors. $\Sigma$ contains the amount of variance in the direction of each principal component, and $V^T$ contains the eigenvectors of $X^TX$. Both U and V form an orthonormal basis for row and column space, respectively, from which our data can be projected to reveal the characteristics of X.

This lossless decomposition approach separates a data stream into components (eigenvalues) ranked by variance. Projection of the multi variate data onto any of the corresponding eigenvectors produces components that each have distinct contributions to the original signal. Typically, noise is a low variance signal. Therefore, by truncating lower-ranked components and retaining higher-ranked components, PCA can selectively isolate the signal from the noise and improve data quality[1,2].

#### 1.2 PCA applied to FLIM

In the case of FLIM imaging with a time-correlated single photon counting (TCSPC) scheme, the detection noise incorporated into the temporal photon count histograms has a Poisson's distribution[3–5]. While every pixel in a FLIM image is inherently a linear combination of multiple lifetimes, this noise disrupts the variance of the actual data and the correlated linearity of the pixels[4,6]. Therefore, the noise profile in FLIM signals, while it is random, does hold some structure in the form of Poisson's noise. To correct for this Poisson profile of photon counting noise and retain the correlated linearity among pixels, we implement a Poisson normalization scheme using the following expression[4]:

$$I^n_{q,m} = \frac{I^r_{q,m}}{\sqrt{\bar{I}^r_m}} \tag{2}$$

where $I^n_{q,m}$ is the normalized intensity at time bin m and pixel q. $I^r_{q,m}$ is the measured pixel intensity, and $\bar{I}^r_m$ is the average measure pixel intensity. This normalization corrects for intensity dependent noise contribution to data variance and is used to reduce noise contributions to higher-ranked principal components. Next, the noise corrected data enters the described PCA workflow. In general, signals produced by fluorophores in FLIM systems are far more structured and less random, which tends towards high directional variance in the signal. In contrast, random noise tends toward lower variance given its random nature and lack of structure. These lower-ranked structures which are dominated by noise are truncated from our matrix decomposition, resulting in a reconstruction of our data with a reduction in noise content.

#### 2 FLIM Data Analysis

As discussed in the main text, there are several strategies to analyze FLIM data. One commonly used approach combines image selection, phasor transformation and thresholding, filtering, and possible image averaging (in this work called Filtered Phasor Analysis). In this work, we analyze data using our newly developed PCA-based approach and compare the results to those obtained using Filtered phasor analysis method.

##### 2.1 Phasor Analysis

The foundation of phasor analysis comes in the form of obtaining a system's phasors, commonly denoted as G and S, through the application of a Fourier transform to time domain fluorescent decay signals. G and S are the real and imaginary components that result from this transformation and are calculated by[7]:

$$G(\omega) = \frac{\int I(t)\cos(\omega t)}{\int I(t)} \tag{3}$$

$$S(\omega) = \frac{\int I(t)\sin(\omega t)}{\int I(t)} \tag{4}$$

here, $I(t)$ is the intensity per pixel in time, and $\omega$ is the angular frequency of the excitation laser.

Every point on the phasor histogram is indicative of a unique lifetime state. The coordinates of single lifetime samples lie on the universal semicircle, while the phasor coordinates of multi-exponential decay samples lie inside of the circle and can be decomposed into a linear combination of single lifetime species on the unit circle. The components of the phasor diagram (G and S) are related to polar coordinate representation of phasor histogram through:

$$|m(\omega)| = \sqrt{(g(\omega)^2 + s(\omega)^2)} \tag{5}$$

$$\varphi = \arctan\left(\frac{s(\omega)}{g(\omega)}\right) \tag{6}$$

where m and φ are the modulation and phase shift of the fluorescent signal for frequency domain measurements[8,9]. For single exponential decays using the definition of s and g, it could be shown that lifetime is related to m and φ through:

$$\tau_{\varphi} = \frac{1}{\omega}\tan(\varphi) \tag{7}$$

$$\tau_{m} = \frac{1}{\omega}\sqrt{\frac{1}{m^2} - 1} \tag{8}$$

where $\tau_m$ and $\tau_\varphi$ are lifetime values related to amplitude and phase modulation. By using standard dyes with known lifetimes, the phasor plot can be calibrated using these equations.

In calculating the phasor components (s and g), the entire signal is incorporated into the integral. Therefore, phasor transformations directly translate noise into s and g. The inclusion of noise increases the error when interpreting the data. This necessitates application of a denoising techniques on FLIM data.

#### 2.2 Denoising Approaches

##### *2.2.1 Thresholding and Background Removal*

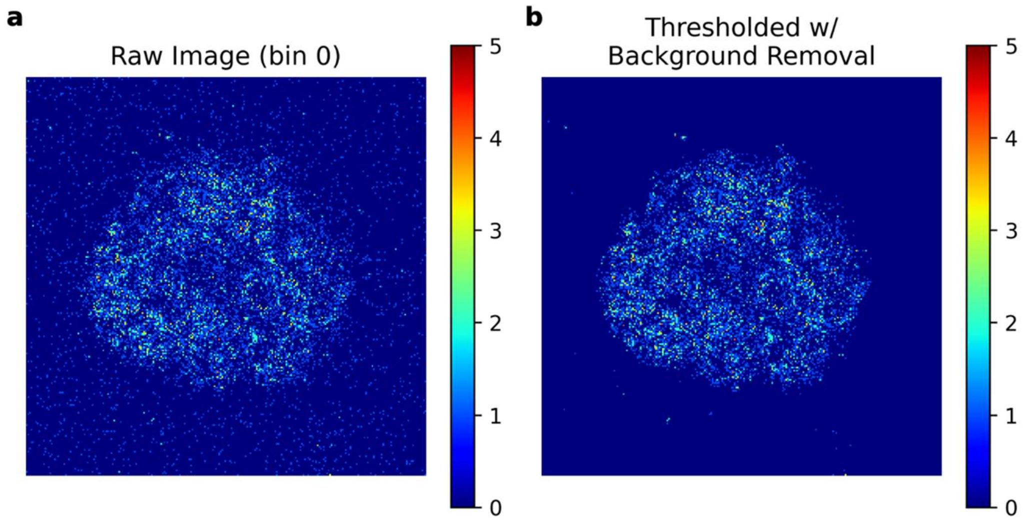

**Figure S1: Effects of thresholding and background removal.** (**a**) Raw image of representative organoid (at t = 0). (**b**) Same image as in (**a**) with intensity thresholding of 8 counts applied.

Intensity thresholding is performed by summing counts in successive time bins (total counts per pixel) and replacing values below a set threshold to zero. Fig.S1 shows a representative thresholded image of FLIM data. In this work, thresholding ranged from 5 to 8 counts and is sample dependent. For our samples, we found a threshold between 5 to 8 was sufficient enough to remove background counts while maintaining fluorophore signal.

Decay background removal scheme was based on a total energy offset captured in the data. Pixel intensities were averaged per time bin. Nearer the last bins, intensities should be zero. When an offset is present, we average the final few bin intensity averages and subtract that DC offset from each time bin.

##### 2.2.2 *Image Selection and Filtering*

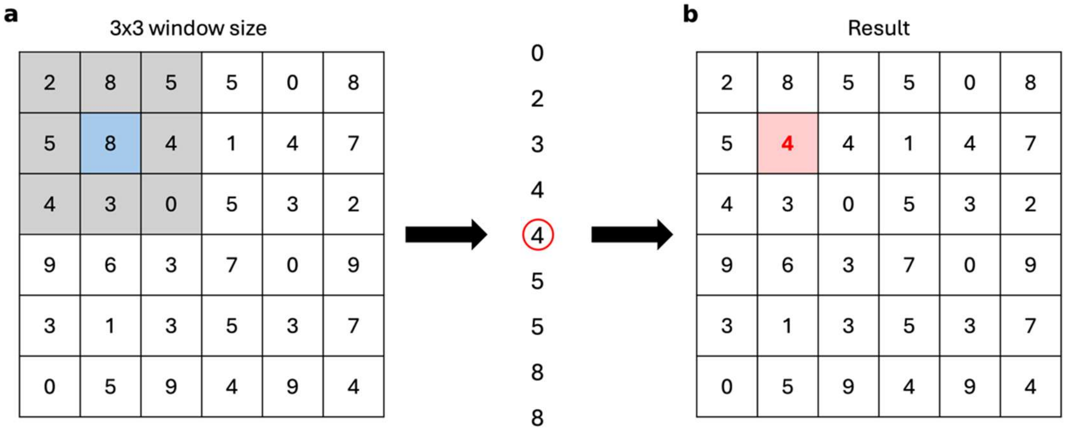

**Figure S2: Example of applying a median filter to a section of data with a 3x3 window size.** (**a**) Data selection (blue) with defined kernel (window) size of 3x3. (**b**) Resulting change in value (red) post median filtering.

For the data used in this work, a manual image selection step was used to remove empty or very weak intensity (~<50 counts average) images. Subsequently, the time domain FLIM data was transformed to phasor domain and thresholding and median filtering was applied. To avoid possible over smoothing in the data we have processed all the data for filtered phasor analysis (method 1 in the main text) with a window size of 3. Fig. S2 shows the process of median filtering.

##### 2.2.3 *NC-PCA*

Prior to PCA, we applied the thresholding and background removal schemes described in in 2.2.1. No image selection or additional filtering was applied beyond NC-PCA. Once images were preprocessed as outlined in section 2.2.1, we applied the noise correction scheme and subsequent decomposition outlined in section 1.1 and 1.2. These noise correction factors are stored for later use in converting our image reconstruction back to its original basis.

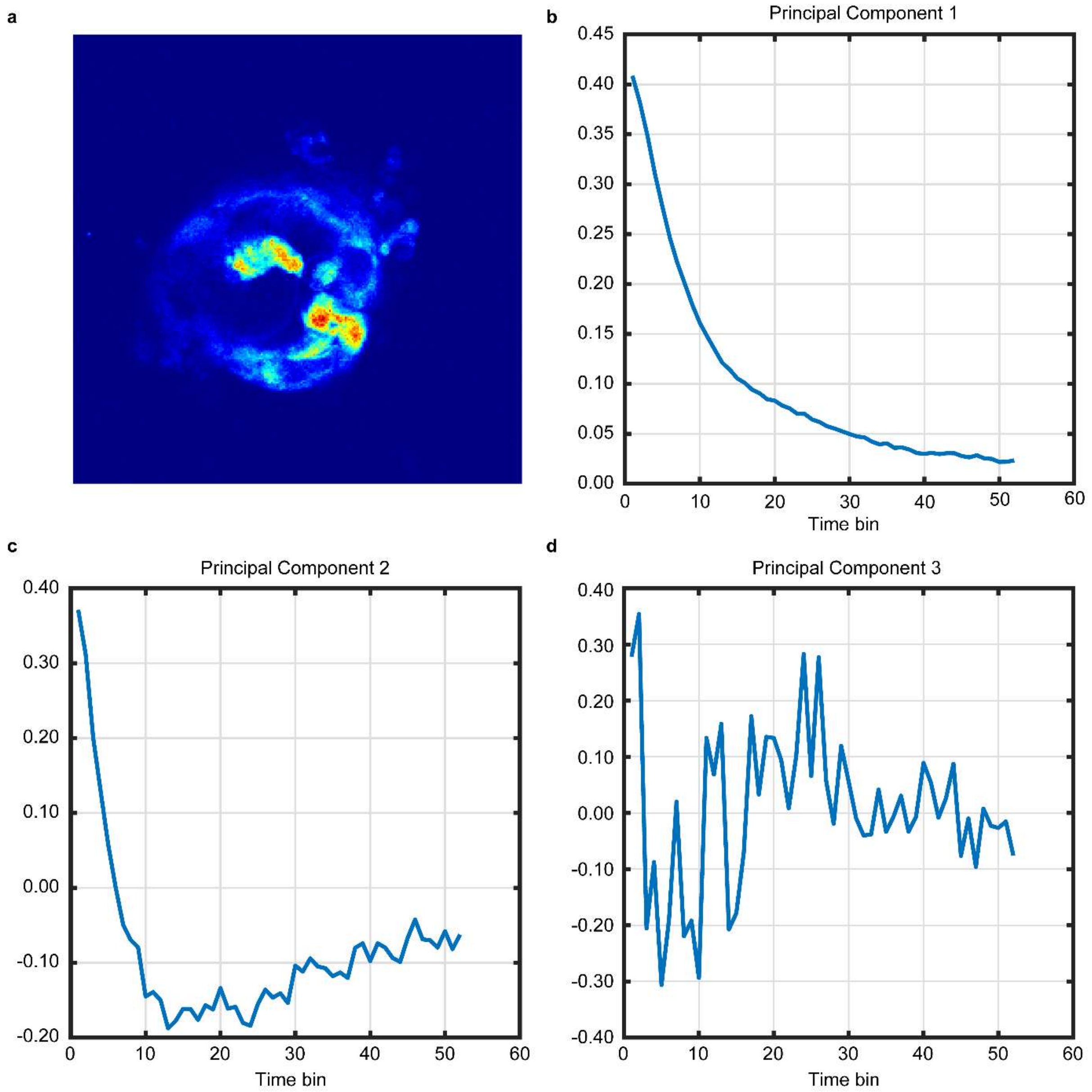

**Figure S3: Representative Example of first 3 principal components in representative dataset using PCA (blue).** (**a**) Representative 0$^{th}$ time-bin FLIM image followed by corresponding (**b**) 1$^{st}$, (**c**) 2$^{nd}$, and (**d**) 3$^{rd}$ principal component.

After we project our dataset onto the selected principal components (Z scores), the data is reconstructed using only the first three projections and finally we scale our results back to its original basis using the stored noise correction factors. Fig. S3 shows a representative image from a patient derived spheroid along with the first three components.

##### 2.3 SNR and MSE quantification

Standard forms of SNR and MSE calculations. SNR in decibels is calculated by

$$P_{signal} = \frac{1}{M}\sum_{i=1}^{M} S[i]^2 \tag{9}$$

$$P_{noise} = \frac{1}{M}\sum_{i=1}^{M} N[i]^2 \tag{10}$$

$$SNR_{dB} = 10\log_{10}\left(\frac{P_{signal}}{P_{noise}}\right) \tag{11}$$

where $\boldsymbol{P_{signal}}$ and $\boldsymbol{P_{noise}}$ is the power of the signal and noise, respectively, for $\boldsymbol{M}$ number of frames (time bins). $\boldsymbol{S}[\boldsymbol{i}]$ and $\boldsymbol{N}[\boldsymbol{i}]$ is the signal and noise at the $\boldsymbol{i^{th}}$ time bin.

MSE is calculated by

$$MSE = \frac{1}{M}\sum_{i=1}^{M}\left(S[i] - \hat{S}[i]\right)^2 \tag{12}$$

where $\boldsymbol{\hat{S}}[\boldsymbol{i}]$ is the ground truth (predicted) value of the signal at the $\boldsymbol{i^{th}}$ time bin.

##### 2.4 Converting G to fraction bound

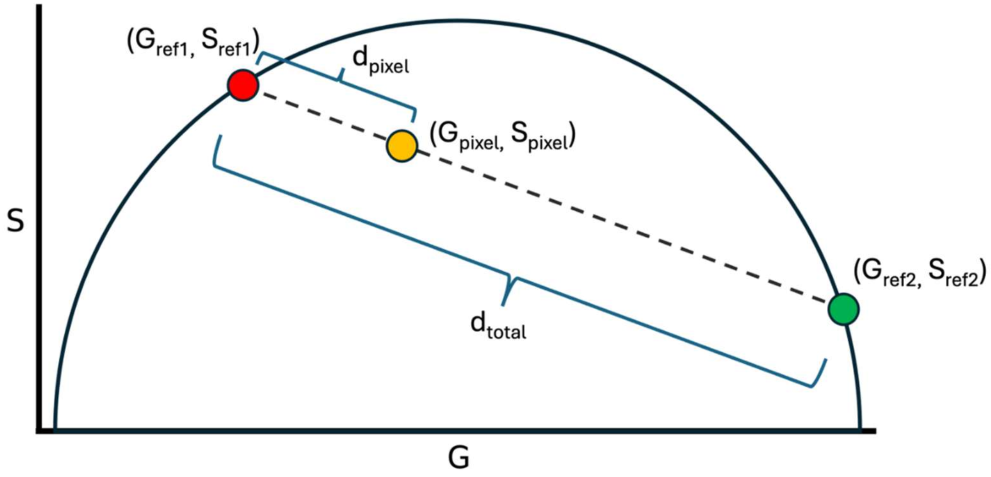

**Figure S4: Diagram of key components for calculating fraction bound for inline data points**

Fraction bound starts by determining the G and S coordinates for two reference lifetimes which lie on the universal semicircle. This is done by the following:

$$G_{ref_{1,2}} = \frac{1}{1+(\omega\tau_{1,2})^2} \tag{13}$$

$$S_{ref_{1,2}} = \frac{\omega\tau}{1+(\omega\tau_{1,2})^2} \tag{14}$$

where $\omega = 2\pi f$ and $f$ is the modulation frequency, $\tau_{1,2}$ represents one of the two reference lifetimes. The distance from each phasor is then calculated using these reference G and S values using a Euclidean distance measurements.

$$d_{pixel_{1,2}} = \sqrt{\left(G_{pixel} - G_{ref_{1,2}}\right)^2 + \left(S_{pixel} - S_{ref_{1,2}}\right)^2} \tag{15}$$

$$d_{total} = \sqrt{\left(G_{ref_2} - G_{ref_1}\right)^2 + \left(S_{ref_2} - S_{ref_1}\right)^2} \tag{16}$$

Here, $d_{pixel_{1,2}}$ is the distance of one phasor to either $\tau_1$ or $\tau_2$, and $d_{total}$ is the total distance between the reference lifetimes on the universal semicircle. As highlighted in Fig.S4 the faction bound is thus calculate by,

$$f_{bound} = \frac{d_{pixel_2}}{d_{total}} \tag{17}$$

It is important to mention that for the experimental data, the G and S coordinates for each pixel might not always be on the line connecting the free and bound phasor coordinates. This can occur due to noise, molecular heterogeneity, or partially bound states[10–13]. To calculate the fraction bound for any point that is not on the line connecting the free and bound NADH end points, we project the vector connecting the coordinates of the data point and the free NADH onto the vector connecting the two end points this process is summarized in Fig. S5. In this case the fraction bound is calculated by:

$$f_{bound} = \frac{projection}{d_{total}} \tag{18}$$

where projection is given by:

$$projection = \frac{\vec{v}_{free\_to\_data} \cdot \vec{v}_{free\_to\_bound}}{d_{total}}$$

(Eq. 19)

where $\vec{v}_{free\_to\_data}$ and $\vec{v}_{free\_to\_bound}$ are the vectors connecting the coordinates of free NADH and the data and free NADH to bound NADH respectively.

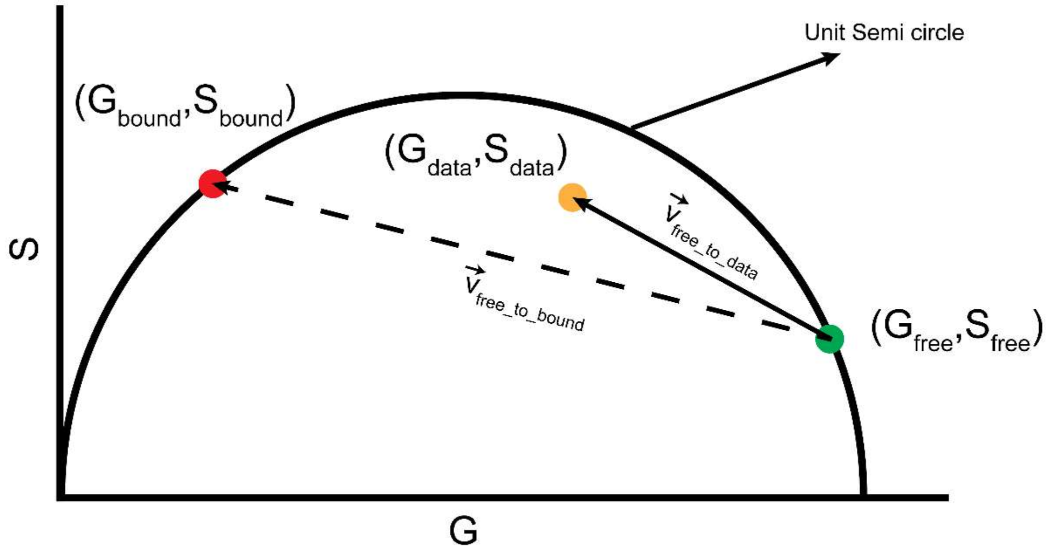

**Figure S5: Diagram of key components for calculating fraction bound for data points outside the reference line**

### 3 PCA analysis code

PCA analysis workflow

Below (Fig. S6) is a simplified workflow for the appropriate application NC-PCA

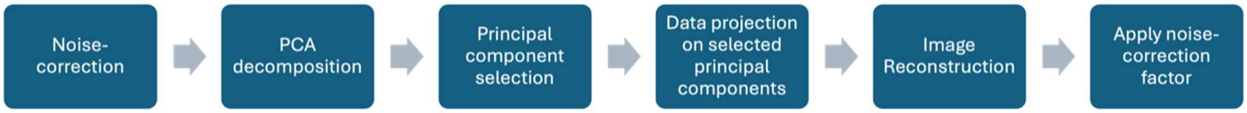

**Figure S6: Simple workflow for NC-PCA**

General workflow as shown in the figure above consists of the initial noise-correction (normalization) step, followed by PCA decomposition and principal component selection. Once principal components are selected, the noise-corrected dataset is projected onto these components, and the signal is reconstructed using the selected principal components and finally the noise-correction factor is applied.

### 4 Synthesized data

#### 4.1 Design and creation of synthesized data

An initial cartoon image of a cell is created in .tiff format. This 540 x 720 pixel image included basic components, including a nucleus, mitochondria, vacuoles, ribosomes, and a cell wall, which were assigned distinct lifetime values. Next to simulate the experimental data we replicated the cartoon image 256 times and multiplied every pixel by the assigned exponential decay function with the designed lifetime value. Chosen lifetimes and their corresponding components in the synthetic data set are presented in Table S1. A range of 1.25 to 2.5 ns were chosen, as they are within a common range of lifetimes for autofluorescent studies of metabolism[14–17]. Finally, for every pixel in FLIM data the noise content is defined using Poisson distribution and pixel counts.

**Table S1: Cartoon cell components and their assigned lifetimes**

| Lifetime (ns) | Component |
|---|---|
| 1.25 | Cell wall |
| 1.90 | Mitochondria, ribosomes |
| 2.50 | Nucleus, vacuoles |

Examples of variations in signal and noise for counts between 200 and 1000 are shown along in figure S7.

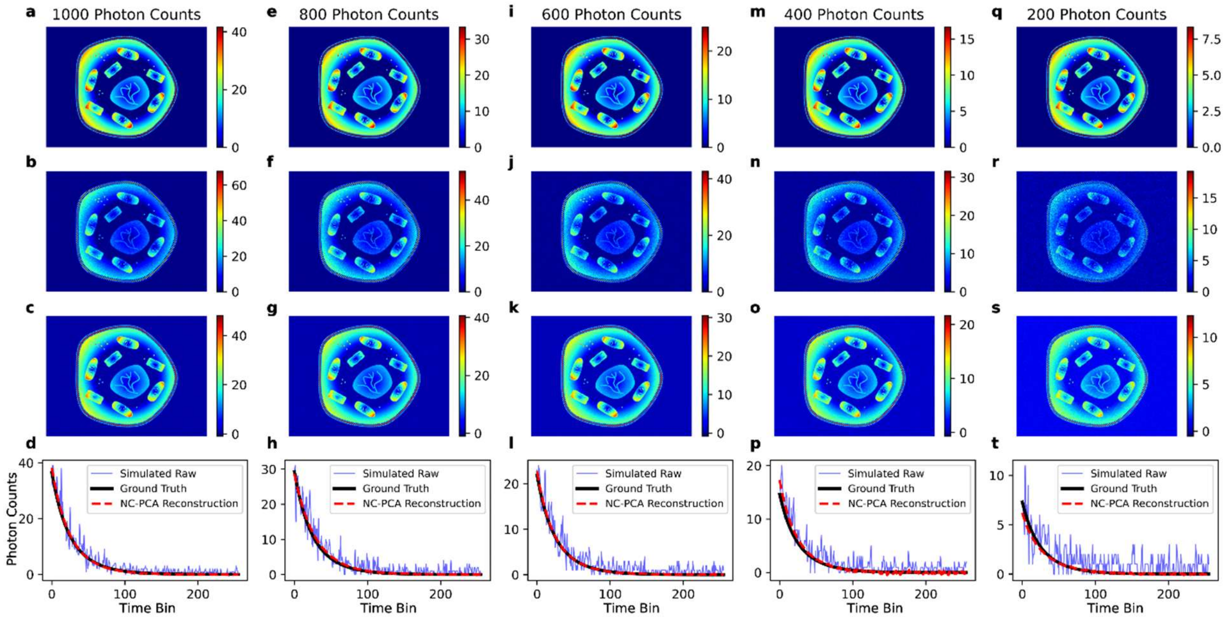

**Figure S7: Examples of noise embedded synthetic data at different total counts per pixel varying from 200 to 1000 counts**. (**a,e,i,m,q**) Ground truth images normalized to 1000, 800, 600, 400, 200 photon counts, respectively. (**b,f,j,n,r**) Images with applied Poissonian noise and (**c,g,k,o,s**) the NC-PCA reconstruction of the synthetic experimental FLIM data. (**d,h,l,p,t**) Representative pixel decay of the ground truth (black), synthetic experimental (raw) data (blue), and the NC-PCA reconstruction (red).

#### 4.2 Error Ratio Trends Over Wide Count Regime

While across all the data sets in this study, NC-PCA generally exceeds median filtering methods however, with the synthesized dataset, we are able to see the efficiency limits for NC-PCA method. To this end, we compared synthetic raw data to NC-PCA reconstructed synthetic images for varying degrees of normalized photon counts. Furthermore, we introduce two metrics: SNR Ratio and MSE Ratio to determine how the SNR and MSE from NC-PCA compare to the synthetic raw datasets. In Fig. S8 we have shown the trends in SNR Ratio and MSE Ratio for count per pixel values ranging from 80-1800.

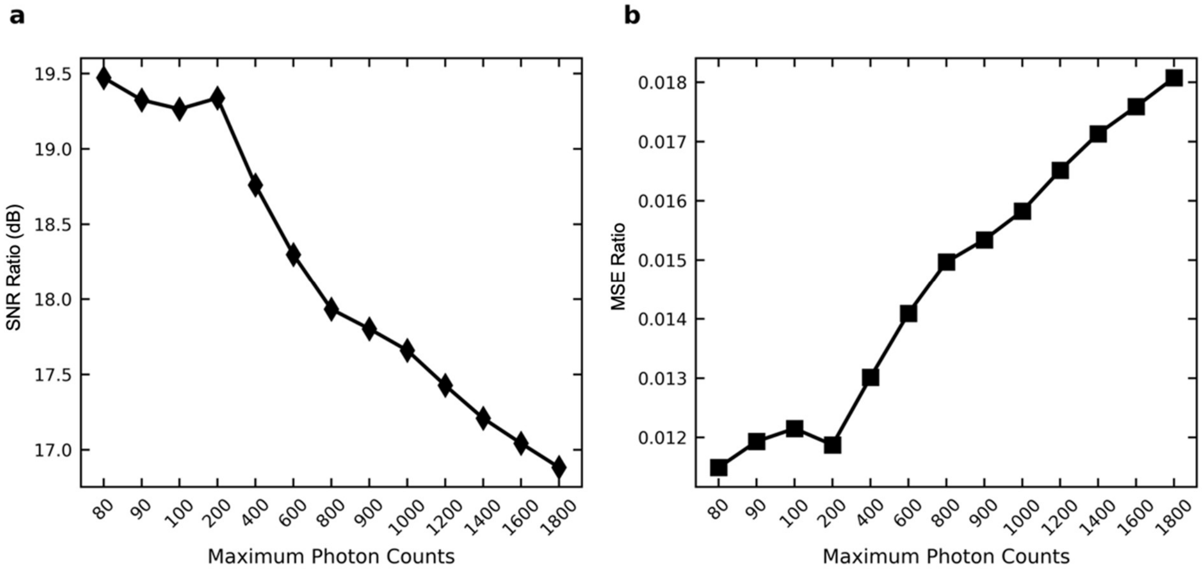

**Figure S8: Calculated SNR and MSE Ratio for varying total photon count range.** (**a**) SNR Ratio and (**b**) MSE Ratio over normalized synthetic FLIM data ranging from 80 to 1800 photon counts

#### 4.3 Calculation of Loss maps

To calculate the map of G and fB values that maintain their original values if phasor histogram averaging is applied, we first calculated the coordinates of average values of G and S. Next on G and fB images we kept the values of G and S that fall within a small radius with respect to average G and S values on the phasor histogram (0.01). The preserved points represent the pixels that aren't affected by histogram averaging.

#### 4.4 NC-PCA validation using experimental FLIM data

When imaging dynamic biological systems using FLIM, the number of acquired frames is often lowered to maintain acceptable image acquisition rates. For example, the number of acquired frames is normally kept in the range of 10-30 frames which allows the minimum total counts per pixel to be greater than 100. By increasing the number of acquired frames to ~100-200 frames, the total photon count increases to 7000-10000, and the accuracy of the data analysis improves. However, this increase in frame number dramatically increases the image acquisition time. Here, we show that NC-PCA allows reliable reconstruction of the time decay signal per pixel in low photon measurements, allowing low frame numbers to be used without compromising SNR.

In the experiment, Coumarin6, a standard fluorophore with a known lifetime in the range of 2.43-2.60 ns[18–20], was imaged with two frame acquisition values: 30 and 100. The decrease in noise during the 100-frame measurement allowed this data to serve as a control (ground truth). Next, we applied a single component exponential fit to every pixel in each image to obtain lifetime values. In addition, NC-PCA was performed on the 30-frame acquisition data, and lifetime values were obtained.

Figure S9a presents a representative temporal intensity decay for the 100-frame ground truth data that is fit to a single exponential decay. Attempts to fit the raw 30-frame data were unsuccessful because they either did not converge for majority of the pixels or resulted in large error due to insufficient signal-to-noise ratios. However, after applying NC-PCA to the 30-frame data, the previously obscured signal became clearly identifiable, and the data could be reliably fit (Figure S9b). Thus, the subsequent discussion focuses on a comparison between the 100-frame ground truth data and the NC-PCA analyzed 30-frame data to highlight the efficacy of NC-PCA in recovering signal from high noise data.

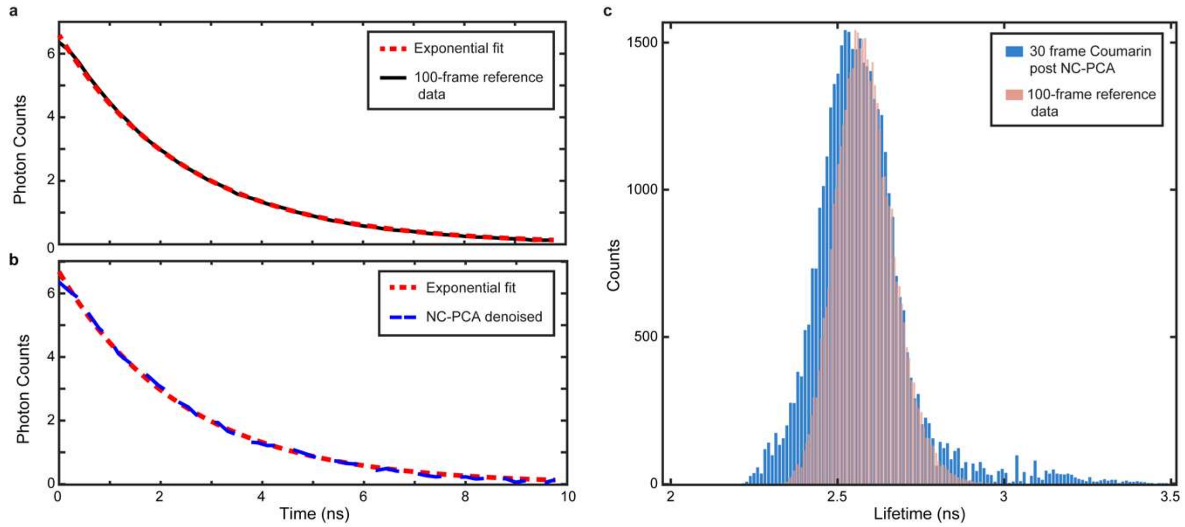

**Figure S9: Comparison of lifetime distribution between the 100-frame ground truth FLIM signal and 30-frame raw data post NC-PCA reconstruction for Coumarin6.** Representative temporal decay FLIM signal overlaid on single exponential fit for (**a**) 100-frame ground truth (**b**) 30-frame raw data post PCA reconstruction. (**c**) comparison of histograms for extracted lifetimes from 100-frame ground truth and post NC-PCA denoising for 30-frame Coumarin6 FLIM images.

Figure S9c shows the lifetime distribution from the ground truth image and from the NC-PCA reconstructed 30-frame FLIM image. The reference histogram has a mean value of 2.58 ns and a Full Width Half Maximum (FWHM) of 0.2 ns, which is consistent with the expected range. While the raw 30-frame data could not be reliably fit, applying NC-PCA to the same data yields a mean lifetime of 2.56 ns and a FWHM of 0.23 ns. These results demonstrate that NC-PCA allows low frame data to achieve a similar fidelity to the high frame data.

#### 5 3D culture preparation, imaging, and analysis

Colorectal cancer tumor resections were received from USC Norris Comprehensive Cancer Center Translational Pathology Core according to Institutional Review Board (Protocol HS-06-00678) approval. Tissues were processed to generate organoids as previously described[21,22]. For Expansion expansion, organoids were maintained in 5% $CO_2$ at $37^0$C in Cultrex Reduced Growth Factor Basement Membrane Extract, Type 2 (BME; R&D Systems, Minneapolis, MN; 3533-001-02) with ADMEM/F12 culture media spiked with the following supplements in Table S2:

**Table S2: CTO media supplements**

| Supplement (solvent if applicable) | Final Concentration | Manufacturer |
|---|---|---|
| Heat inactivated fetal bovine serum | 10% | GeminiBio; 100-500 |
| Penicillin : Streptomycin | 5% | GeminiBio; 400-109-100 |
| HEPES | 1% | Thermofisher Scientific; 15630080 |
| GlutaMAX | 1% | Thermofisher Scientific; 35050-061 |
| B27 supplement | 2% | Thermofisher Scientific; 17504-001 |
| N2 supplement | 1% | Thermofisher Scientific; 17502048 |
| N-acetylcysteine (water) | 1 mM | Sigma Aldrich, A9165 |
| Nicotinamide (water) | 10 mM | Sigma Aldrich, N0636 |
| Noggin (0.1% BSA in PBS) | 100 ng/mL | Peprotech, 120-10C |
| EGF (1% BSA in PBS) | 50 ng/mL | Thermofisher Scientific; PHG0313 |
| SB202190 (DMSO) | 10 µM | Sigma Aldrich, S7067 |
| A-83-01 (DMSO) | 500 nM | Millipore, 616454-2MG |

For FLIM experiments organoids were plated as previous described[23]. Briefly, organoids were dissociated into single cells by dissolving BME with Gentle Cell Dissociation Reagent (STEMCELL Technology, Cambridge, MA; 07174) at $4^0$C with rocking for 30-40 minutes. A P-1000 pipette tip was used to break up the organoid fragments before centrifuging at 300xg for 5 minutes and replacing supernatant with 1:1 PBS:TrypLE (Thermofisher Scientific, Waltham, MA; 12605028 spiked with Y-27632 (STEMCELL Technology, Cambridge, MA; 72302) at 1:1000 dilution. The cell suspension was passed through a 40µm cell strainer before isolating the cell pellet through centrifugation and resuspending in CTO media. 1000 cells/well were plated in a glass-bottom 96 well plate (Mattek, Ashland, MA; P96G-1.5-5-F) and grown in CTO media for 7 days before treatment.

Five different treatment strategies were studied in a pair of experiments, as outlined in Table S3. Treatments were applied 72 hours before the first imaging sessions, and imaging was performed at 6 and 72 hours. All therapeutic concentrations were created using serial dilution.

The second study was investigating the role of cancer associated fibroblasts (CAF) in therapeutic efficacy. The following series of FLIM measurements were performed: no treatment, addition of CAF to culture media, treatment with 3-BP, culture with CAF and subsequent treatment with 3BP (3BP-CAF).

**Table S3: Therapeutic treatment conditions**

| Treatment | Vendor | Catalog # | Concentration Range |
|---|---|---|---|
| **Study 1** | | | |
| Staurosporine (ST) | Sigma-Aldrich | 569396 | 0, 0.01μM, 0.1 μM, 1μM, 10 μM |
| 5-fluorouracil (5-FU) | Selleck Chemicals | S1209 | 0, 0.1 μM ,1μM, 10 μM, 50 μM |
| 7-ethyl-10-hydroxycamtothecin (SN-38) | Sigma-Aldrich | H0165 | 0, 0.01μM, 0.1 μM, 1μM, 10 μM |
| Cetuximab (CTX) | MedChemExpress | HY-P9905 | 0, 0.01μM, 0.1 μM, 1μM, 10 μM |
| **Study 2** | | | |
| 3-bromopyruvate (3-BP) | Sigma-Aldrich | 16490 | 0, 25 μM, 50 μM, 80 μM, 100 μM |

#### 6 Metabolic effects

To confirm the universality of the NC-PCA method, a secondary analysis was performed on the FLIM studies summarized in Table S3. Notably, this pair of investigations spanned several different therapeutics and were performed on two different microscopes.

#### 6.1 Applying NC-PCA to investigate different therapeutic agents

To show NC-PCA can effectively improve the phasor histogram resolution and thereby unveil effect of drugs on metabolic activity of organoids, we studied the effect of standard chemotherapy drugs for colorectal cancer on PDTOs. For this study, SN38, 5-FU, and CTX were used and the PDTOs were imaged at 6 hours and 72 hours. In addition, ST was used as positive control. Fig. S10-S13 summarize the results.

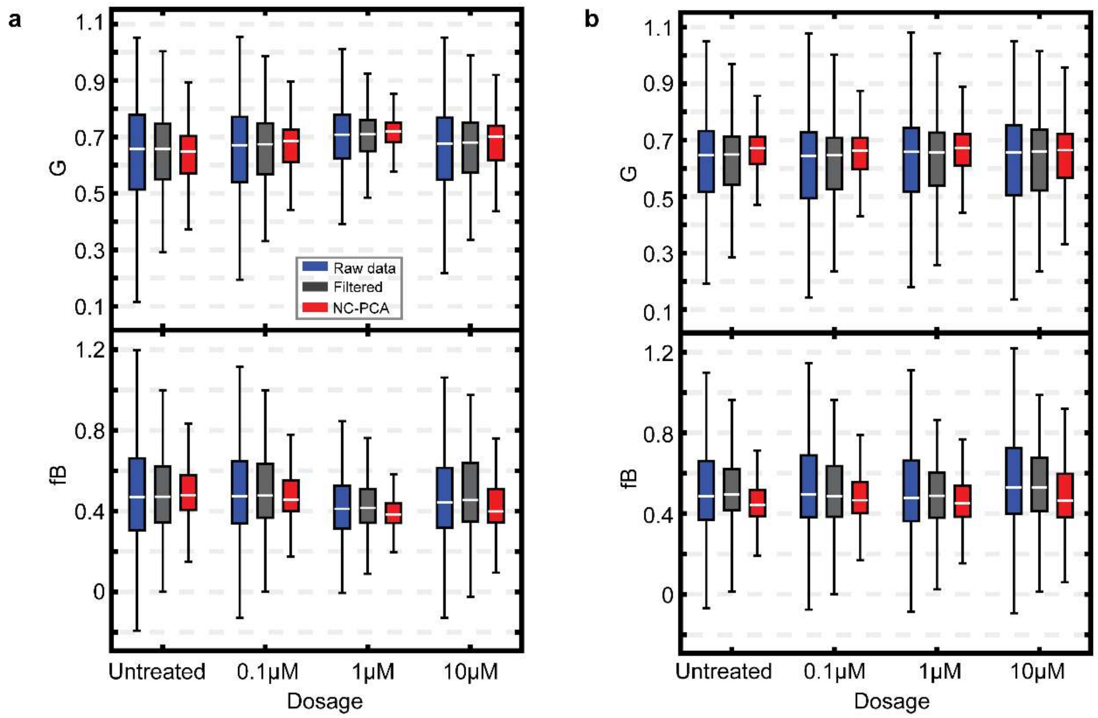

**Figure S10: Calculated boxplots for G and fraction bound (fB) for 5-FU.** (**a**) 6 and (**b**) 72 hours. All the calculations were done with an applied threshold of 8 and a median filter with a window size of 3.

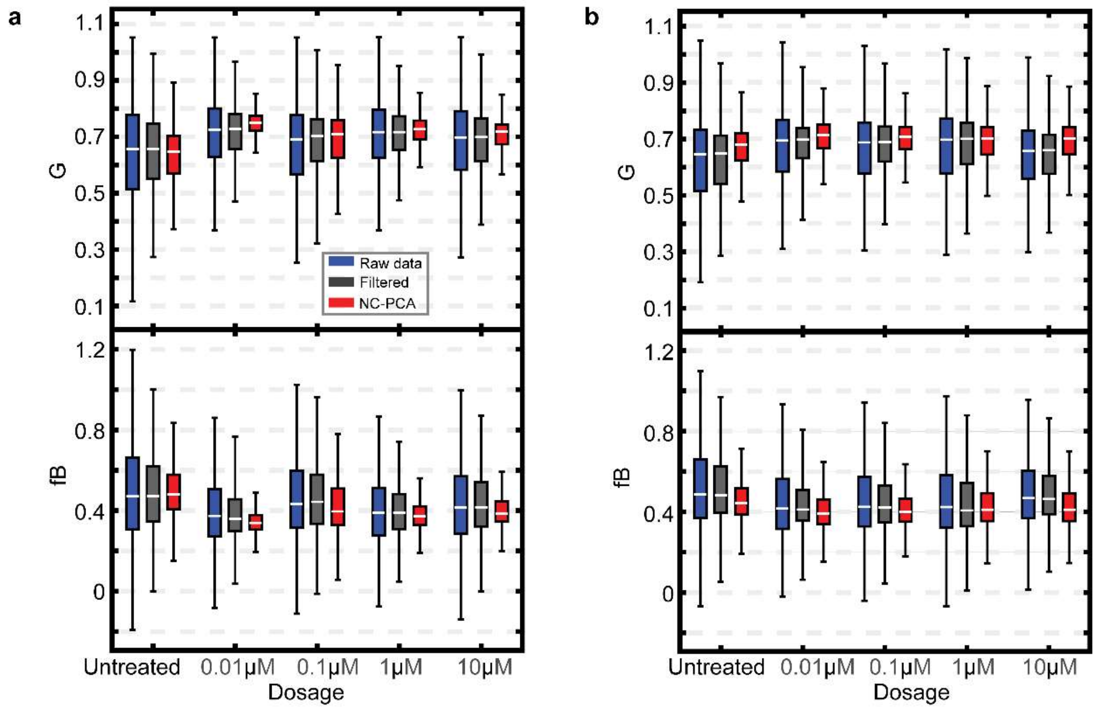

**Figure S11: Calculated boxplots for G and fraction bound (fB) for CTX.** (**a**) 6 and (**b**) 72 hours. All the calculations were done with an applied threshold of 8 and a median filter with a window size of 3.

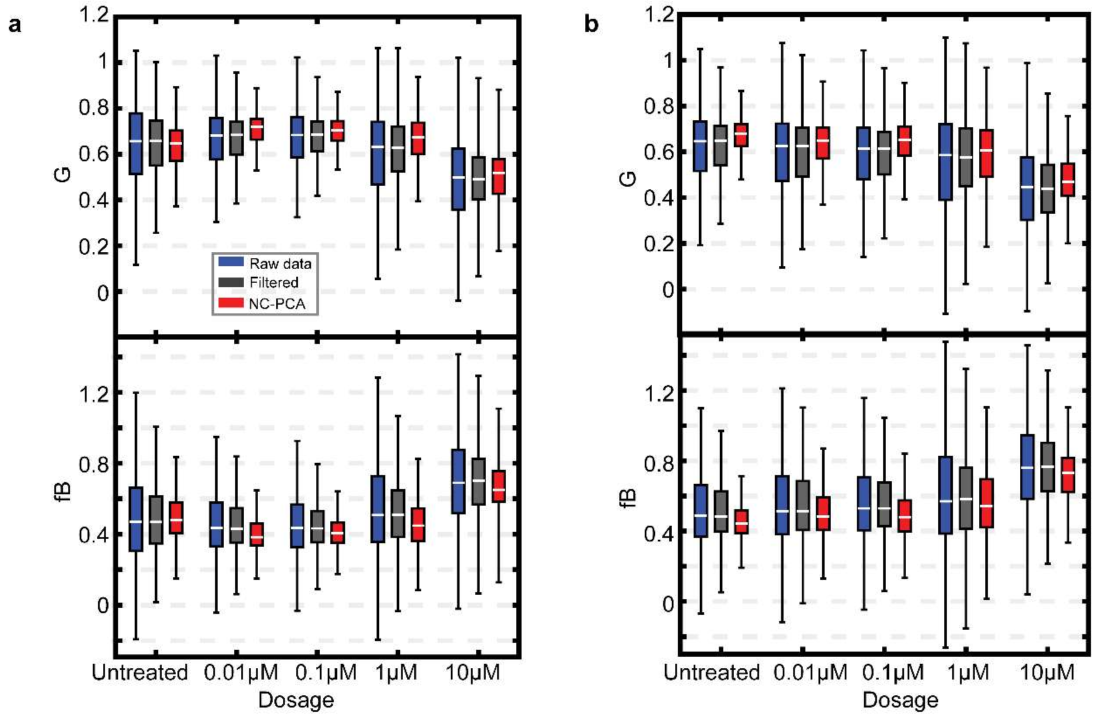

**Figure S12: Calculated boxplots for G and fraction bound (fB) for SN38.** (**a**) 6 and (**b**) 72 hours. All the calculations were done with an applied threshold of 8 and a median filter with a window size of 3.

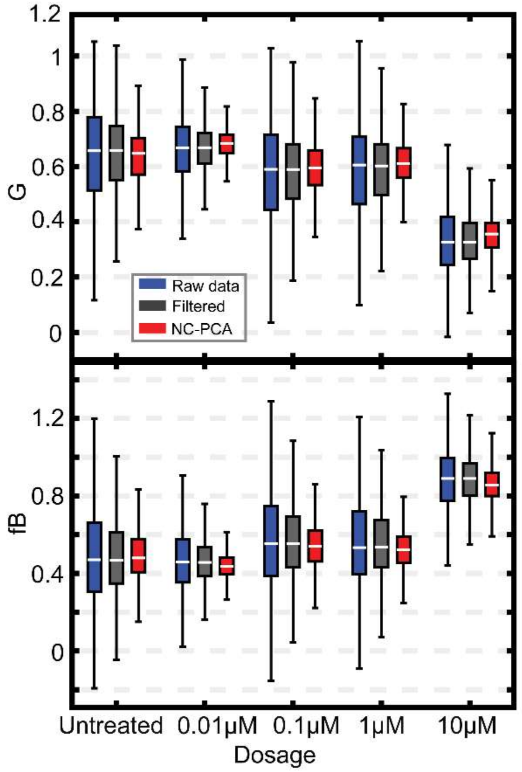

**Figure S13: Calculated boxplots for G and fraction bound (fB) for ST.** (**a**) 6 and (**b**) 72 hours. All the calculations were done with an applied threshold of 8 and a median filter with a window size of 3.

#### 6.2 Using NC-PCA with different imaging systems

To further confirm the universality of the analysis capability, we apply it to images taken with a different microscope, the Olympus FV3000 system connected to an A320 FastFLIM FLIMbox (ISS Inc., Champaign, IL). Moreover, we studied a different series of therapeutic treatments. The organoid samples were treated with one of the following therapeutics 3BP, 3BP with cancer associated fibroblast (3BP-CAF), 5FU, or SN38 at varying concentrations. We have summarized the results in Fig. S14.

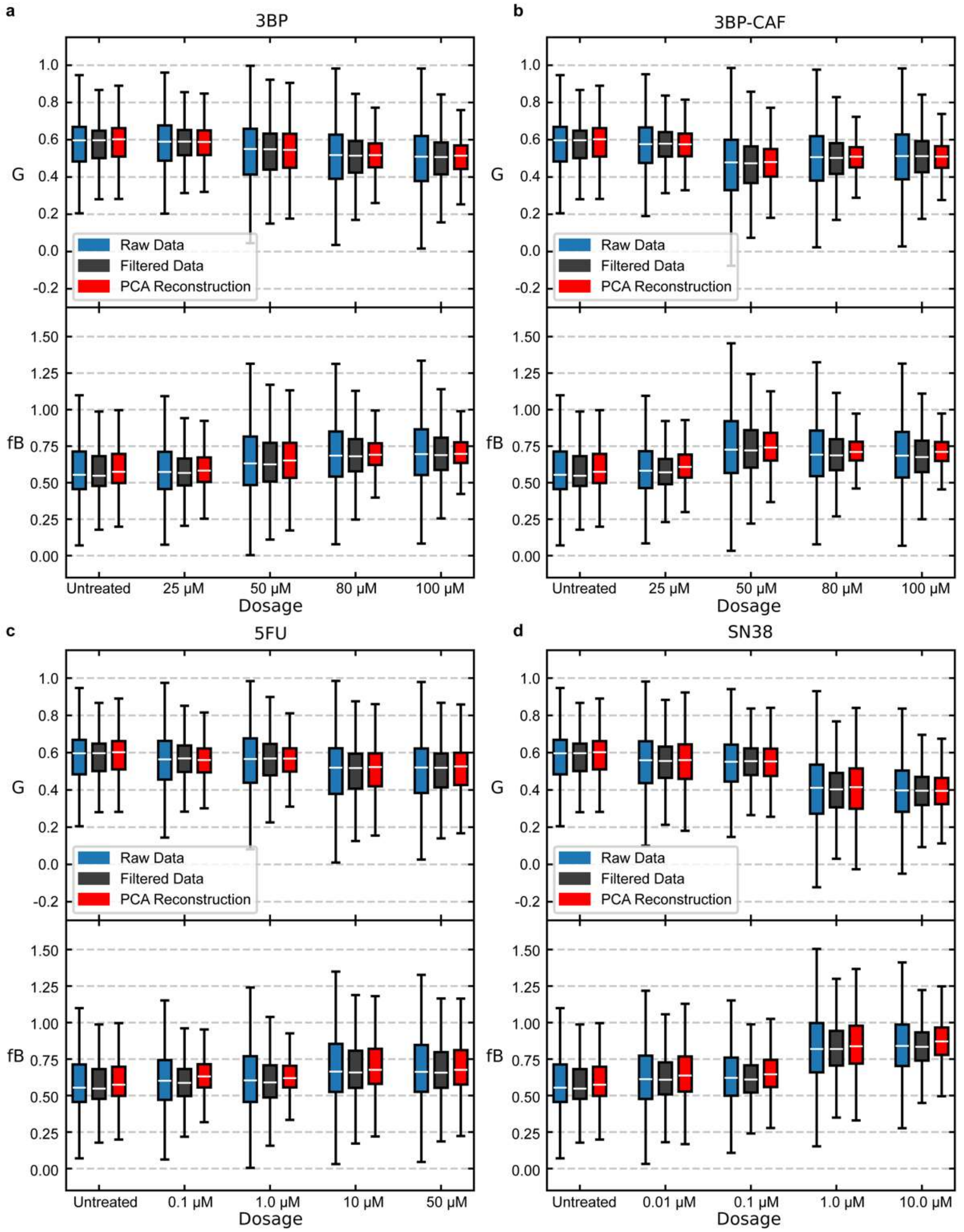

**Figure S14: Additional G and fB analysis.** (a) 3BP, (b) 3BP-CAF, (c) 5FU, and (d) SN38 using a separate FLIM system to what was presented in the main text. Analysis was done with an applied intensity threshold of 8 and median filter window size of 3. NC-PCA maintained the first three primary principal components for projection and reconstruction.

As referenced in the main text and shown via the SNR Ratio and MSE Ratio, systems with higher photon counts tend to produce results with higher SNR. The cost benefit analysis of PCA when applied to high SNR systems is diminishing. While for the majority of drugs and selected treatment we see improvement after NC-PCA, we observe the results previously studied when no longer in a low count regime. Regardless, NC-PCA often matches and exceeds that of the filtered data without the risk of smoothing errors.

#### 7 Experimental metadata

All metadata for all data sets used in this work is shown below for transparency. When the data sets are included in a figure in the paper, the figure is indicated. The experiments are presented in the order in which they appear in the main text and SI.

**Table S1: Image and Experimental hardware specifications.**

| **Experimental/Sample** | |
|---|---|
| Experimenter Name | Emma Fong, Soheil Soltani, Jack Paulson |
| Experiment Description | imaging and metabolic activity monitoring of PDTOs under different treatments |
| Experiment Date(s) | 2021-06-13 through 2021-06-20, 2021-06-28 through 2021-07-04, 2022-04-27 through 2022-05-03, 2022-05-17 through 2022-05-23, 2023-04-05 through 2023-04-11 |
| Sample Description | Patient Derived Colorectal Cancer tumor organoids |
| Medium | Complete feeding medium, made as previously reported (1) |
| Temperature | $37^0$C |
| CO2 | 5% |
| **Microscope hardware specifications** | |
| Microscope manufacturer(s) | Zeiss |
| Microscope model(s) | 780 AxioObserver inverted |
| Objective manufacturer | Zeiss |
| Objective model | Plan-Apochromat M27 |

| | |
|---|---|
| Magnification/NA | 20x/0.8NA |
| External Detectors | Hamamatsu R928 PMT |
| FLIM hardware | ISS A320 FastFLIM |
| **Image acquisition settings** | |
| Illumination type | Ti:Sapphire 2-photon laser |
| Channel name | External |
| Laser manufacturer | Coherent |
| Laser model | Chameleon Ultra II |
| Pixel dwell time | 12.6 µs/pixel |
| Image size | 256x256 |
| Number of frames averaged | 4 |
